# Antidepressant mechanism and treatment response define distinct hippocampal-amygdala circuit biomarkers during emotional memory in humans

**DOI:** 10.64898/2026.06.01.726320

**Authors:** Madelyn Castro, Hannah Ballard, Lorena Ferguson, Stephanie L. Leal

**Affiliations:** Department of Psychological Sciences, Rice University, 6100 Main St. Houston TX 77005; Department of Integrative Biology & Physiology, University of California, Los Angeles, CA

**Keywords:** depression, antidepressants, memory, hippocampus, amygdala, pattern separation

## Abstract

Antidepressant efficacy varies widely, yet the circuit-level mechanisms that distinguish treatment responders from non-responders remain poorly understood in humans. Here, we used high-resolution neuroimaging of hippocampal-amygdala networks during an emotional mnemonic discrimination task that taxes hippocampal pattern separation to examine how antidepressant mechanism of action and perceived treatment response shape emotional memory circuitry (N = 117). Participants included individuals taking single-action antidepressants (selective serotonin reuptake inhibitors), multi-action antidepressants (serotonin-norepinephrine reuptake inhibitors, norepinephrine-dopamine reuptake inhibitors, or polypharmacy), and unmedicated controls matched on current depression severity. Antidepressant mechanism and treatment response were associated with distinct patterns of activity in hippocampal subfields (dentate gyrus (DG)/CA3 and CA1) and amygdala subnuclei, including the basolateral amygdala (BLA) and central amygdala (CEA), during emotional mnemonic discrimination. Among non-responders, the relative balance of hippocampal activity differed by antidepressant mechanism: those taking single-action antidepressants showed greater DG/CA3 than CA1 activity, whereas those taking multi-action antidepressants showed the opposite pattern. This suggests mechanistically specific differences in hippocampal computations associated with ineffective treatment. These effects were localized to the anterior hippocampus, with no significant effects observed in posterior regions. In contrast, responders exhibited stronger DG/CA3-BLA coactivation during negative mnemonic discrimination, a pattern absent in non-responders and unmedicated controls. Antidepressant-associated differences in amygdala subnuclei activity persisted beyond current symptom severity, suggesting medication-related modulation of emotional memory circuits rather than effects driven solely by depression severity. These findings provide evidence in humans that antidepressant use is associated with altered hippocampal-amygdala circuitry in a manner that depends on both pharmacological mechanism and treatment efficacy. Identifying circuit-level signatures of treatment response may inform mechanistically guided approaches to antidepressant selection and monitoring.

## Introduction

Antidepressants are the primary pharmacological treatment for depression, yet treatment response remains highly variable, with approximately half of patients failing to show meaningful improvement following initial trials (1,2). This variability highlights a critical gap in understanding how antidepressants influence the neural circuits underlying depressive symptoms. Traditional monoamine-based models, particularly the serotonin hypothesis, do not fully account for delayed onset or limited efficacy, motivating growing interest in circuit- and plasticity-based mechanisms of antidepressant action (3).

Beyond mood symptoms, depression is associated with pronounced cognitive dysfunction, particularly deficits in episodic memory and a bias toward remembering negative information (4–13). These impairments are closely linked to dysfunction within medial temporal lobe (MTL) circuitry, including hippocampal subfields and amygdala subnuclei that support emotional memory (14). Notably, the anterior hippocampus, which is more strongly interconnected with the amygdala and supports emotional memory processing, is thought to be particularly relevant for affective dysfunction in depression, whereas the posterior hippocampus is more involved in spatial and contextual representations (15,16). Structural and functional alterations of the hippocampus and amygdala are well documented in depression (17–19), and animal models indicate that antidepressants can modulate hippocampal plasticity, dentate gyrus (DG) neurogenesis, and amygdala reactivity (20,21). However, the extent to which antidepressant treatment alters these circuit-level computations in humans remains unclear (22,23).

One promising framework for characterizing memory dysfunction in depression is hippocampal pattern separation, a computation that supports discrimination among overlapping experiences and depends critically on the DG/CA3 (8,24). Pattern separation can be assessed in humans using mnemonic discrimination tasks (MDT) that include similar lure stimuli (25,26). Prior work has shown that depressive symptoms are associated with altered emotional mnemonic discrimination and aberrant engagement of hippocampal and amygdala subregions, particularly the DG/CA3 and basolateral amygdala (BLA) during negative memory processing (21,25). Importantly, manipulations that enhance hippocampal plasticity and neurogenesis improve pattern separation, suggesting that this computation may be sensitive to antidepressant effects (27).

Furthermore, antidepressants differ substantially in their mechanisms of action (28). Selective serotonin reuptake inhibitors (SSRIs) target a single neurotransmitter system (serotonin), whereas serotonin-norepinephrine reuptake inhibitors (SNRIs), norepinephrine-dopamine reuptake inhibitors (NDRIs), and polypharmacy (POLY; i.e. taking more than one antidepressant) engage multiple neuromodulatory systems (serotonin, norepinephrine, and dopamine) that differentially influence hippocampal-amygdala circuitry (29–31). In addition, antidepressant treatment has been shown to increase neural progenitor cell number in the anterior human DG, suggesting a regionally specific effect on hippocampal plasticity that may be particularly relevant for emotional memory processes (32). Whether these mechanistic differences translate to distinct neural profiles during emotional memory processing and whether such profiles distinguish treatment responders from non-responders remains largely unexplored (22,23).

In the present study, we used high-resolution functional neuroimaging during an emotional MDT to examine how antidepressant mechanism (single-action vs. multi-action) and perceived treatment response (responder vs. non-responder) are associated with hippocampal and amygdala subfield function. We stratified the hippocampal subregions by anterior and posterior axis given prior work showing greater effects of depression and antidepressants on anterior hippocampus. We further compared medicated participants to unmedicated individuals matched on current depression severity to isolate antidepressant-associated effects beyond symptom level. We hypothesized that treatment response and antidepressant mechanism would be associated with distinct patterns of hippocampal subfield recruitment and hippocampal-amygdala coactivation, particularly during negative emotional mnemonic discrimination (25,32,33).

## Methods and Materials

### Participants

Participants (N = 121; ages 18-35) were recruited from the Houston community and completed a single study visit at the Baylor College of Medicine Core for Advanced Magnetic Resonance Imaging (CAMRI). Participants were screened for MRI safety and current medication use. Eligible participants were either (i) currently taking an antidepressant (SSRI, SNRI, NDRI, or more than one antidepressant) for at least one month, (ii) unmedicated individuals experiencing depressive symptoms, or (iii) unmedicated individuals with minimal depressive symptoms. Participants were excluded if they were taking anxiolytics, antipsychotics, mood stabilizers, stimulants, or benzodiazepines, or if they had MRI contraindications. To minimize the influence of extreme values while preserving statistical power, outlier screening was conducted separately for each behavioral and neuroimaging measure. Participants with widespread outlier values or imaging-quality concerns (defined as affecting more than 25% of measures) were excluded entirely from the study (N = 4 with ±5 standard deviations), resulting in a final sample of N = 117. For participants with isolated outlier observations (affecting fewer than 25% of measures), only the specific values exceeding ±5 standard deviations from the sample mean were removed, and the participant was retained for all unaffected analyses. As a result, the sample size may vary slightly across analyses. Reported degrees of freedom reflect the participants included in each specific analysis. All participants provided written informed consent in accordance with the Rice University Institutional Review Board. Participants were compensated with either a $50 Amazon gift card ($25/hr) or 2 Psychology course credits (1 credit/hr) upon completion of the study. To achieve sufficient power to examine group differences, we performed *a priori* power analyses in G*Power 3.1 (34) for a mixed ANOVA design (within-between interaction) across groups (Single vs. Multi; R vs. NR) using a medium effect size (f = 0.25, α = .05, power = .80), which indicated a required sample of 28 participants per group.

### Antidepressant mechanism and treatment response

Participants taking antidepressants were grouped based on pharmacological mechanism as either single-action targeting only serotonin (SSRIs; N = 37) or multi-action targeting combinations of norepinephrine, dopamine, and serotonin (SNRI, NDRI, POLY; N = 32). The single-action group consisted exclusively of participants taking SSRIs, whereas the multi-action group included participants taking SNRIs, NDRIs, or combinations of antidepressants. Thus, group classification reflected the number of monoaminergic systems targeted rather than the number of medications prescribed. Because the study was not powered to examine individual medications separately, analyses focused on broad mechanism-level distinctions reflecting single versus multi-neurotransmitter modulation.

Treatment response was operationalized using retrospective change in depressive symptoms, a commonly used approach when pre-treatment clinical measures are unavailable in cross-sectional samples of treated individuals (35,36). Current depressive symptoms were assessed using the Beck Depression Inventory-II (BDI-II) (37). Retrospective depressive symptoms prior to initiating antidepressant treatment were assessed using modified instructions on the BDI-II, asking participants to report symptom severity before starting medication. A retrospective change score was calculated as the percent reduction from baseline. Participants reporting ≥50% symptom improvement were classified as ‘responders’(R), whereas those reporting <50% improvement were classified as ‘non-responders’(NR), consistent with clinical definitions of antidepressant response (38,39). Participant demographics can be found in Table 1. Importantly, R and NR did not differ in retrospective baseline depression severity, indicating comparable pre-treatment symptom levels. While retrospective reports may be influenced by recall bias, this approach allowed us to capture perceived treatment efficacy in a naturalistic sample and has demonstrated sensitivity to antidepressant-related differences in emotional memory in prior work (32). This strategy also mitigates regression-to-the-mean concerns associated with relying solely on current symptom severity.

**Table 1.** Participant Demographics.

| Variable | Responders<br>Mean (*SD*) | Non-Responders<br>Mean (*SD*) | Single-Action<br>Mean (*SD*) | Multi-Action<br>Mean (*SD*) |
| --- | --- | --- | --- | --- |
| *N* | 47 | 22 | 37 | 32 |
| Subgroup | Single: 20<br>Multi: 27 | Single: 17<br>Multi: 5 | R: 20<br>NR: 17 | R: 27<br>NR: 5 |
| Age | 23.5 (5.0) | 22.4 (3.2) | 23 (4.9) | 23.3 (4.0) |
| Sex | 29F, 10M,<br>6N, 2T | 17F, 3M, 2N | 28F, 5M, 4N | 18F, 8M,<br>4N, 2T |
| Race |  |  |  |  |
| - White | 70% | 59% | 59% | 75% |
| - Asian/Asian American | 26% | 32% | 35% | 19% |
| - Black/African American | 9% | 18% | 10% | 13% |
| - American Indian/Alaska<br>Native/Indigenous | 0% | 0% | 3% | 3% |
| - Arab/Middle<br>Eastern/North African | 0% | 0% | 0% | 0% |
| - Native Hawaiian/Pacific<br>Islander | 0% | 0% | 0% | 0% |
| - Multiracial | 6% | 9% | 8% | 6% |
| - Other | 2% | 0% | 3% | 0% |
| Ethnicity |  |  |  |  |
| - Hispanic/Latinx/a/o | 19% | 23% | 22% | 19% |
| Education (years completed) | 15.4 (2.6) | 15.5 (2.4) | 15.3 (2.4) | 15.7 (2.6) |
| **Questionnaires** |  |  |  |  |
| BDI-II (current)*<sup>+</sup> | 9.6 (6.4) | 24.7 (8.2) | 16.6 (10.2) | 11.8 (9.2) |
| BDI-II (retrospective) | 37.1 (10.2) | 33.7 (10.6) | 35.8 (9.3) | 36.3 (11.6) |
| BAI (current)* | 12.0 (8.8) | 19.2 (11.6) | 15.5 (10.7) | 13.0 (9.8) |
| BAI (retrospective) | 22.8 (12.4) | 24.6 (14.9) | 23.6 (12.7) | 23.1 (13.8) |
| Retrospective Change in<br>Depression | 73.4 (14.7) | 24.9 (19.3) | 50.9 (26.9) | 66.2 (27.2) |
| PHQ-9 | 5.3 (3.3) | 11.0 (5.7) | 7.7 (5.6) | 6.5 (4.2) |
| MDQ | 5.1 (3.8) | 4.8 (3.2) | 4.9 (3.8) | 5.2 (3.4) |
| ASRMS | 2.9 (3.1) | 2.8 (2.5) | 2.7 (2.8) | 3.1 (3.0) |
| Duration of antidepressant use |  |  |  |  |
| - 1 Month | 6% | 0% | 0% | 9% |
| - 2-4 months | 9% | 9% | 11% | 6% |
| - 4-6 months | 4% | 18% | 8% | 9% |
| - 6-11 months | 13% | 5% | 11% | 9% |
| - 1-2 years | 23% | 18% | 35% | 9% |
| - 2-5 years | 19% | 32% | 19% | 25% |
| - 5-10 years | 19% | 14% | 11% | 25% |
| - More than 10 years | 6% | 5% | 5% | 6% |
| Number of switches<sup>+</sup> | 1.0 (1.5) | 0.9 (0.149) | 0.6 (0.8) | 1.4 (1.7) |
| **Neuropsychiatric Measures** |  |  |  |  |
| Perceived stress scale (past month)* | 15.6 (6.8) | 22.4 (4.8) | 19.0 (7.2) | 16.3 (6.5) |
| Perceived stress scale (current)*<sup>+</sup> | 2.8 (2.5) | 6.4 (2.4) | 4.6 (3.0) | 3.2 (2.7) |
| Hours of sleep (last night) | 7.8 (3.6) | 6.4 (1.9) | 7.2 (2.7) | 7.5 (3.8) |
| Hours or sleep (past month) | 8.8 (10.4) | 6.9 (1.2) | 9.2 (11.7) | 7.1 (1.1) |
| ASRS | 11.1 (4.1) | 12.2 (4.1) | 11.2 (4.0) | 11.7 (4.2) |
| Y-BOCS | 7.8 (6.7) | 8.8 (5.2) | 7.7 (5.6) | 8.6 (7.0) |
| **Behavioral Task Accuracy** |  |  |  |  |
| Negative lure CR | 0.69 (0.11) | 0.71 (0.13) | 0.69 (0.13) | 0.71 (0.11) |
| Neutral lure CR | 0.77 (0.11) | 0.79 (0.11) | 0.75 (0.11) | 0.80 (0.11) |
| Positive lure CR | 0.66 (0.14) | 0.71 (0.13) | 0.67 (0.14) | 0.68 (0.13) |
| Negative lure FA | 0.31 (0.11) | 0.29 (0.13) | 0.31 (0.13) | 0.29 (0.11) |
| Neutral lure FA | 0.23 (0.11) | 0.21 (0.11) | 0.25 (0.11) | 0.20 (0.11) |
| Positive lure FA | 0.34 (0.14) | 0.29 (0.13) | 0.33 (0.14) | 0.32 (0.13) |

**Key:** ***BDI-II:** Beck Depressive Inventory II; **BAI:** Beck Anxiety Inventory; **RCD:** Retrospective Change in Depression; **PHQ-9:** Patient Health Questionnaire; **MDQ:** Mood Disorder Questionnaire; **ASRS:** Altman Self-Rating Mania Scale; **ASRS:** The Adult ADHD Self-Report Scale (>14 suggests presence of ADHD); **Y-BOCS:** Yale-Brown Obsessive Compulsive Scale (0-40, larger total scores indicate greater symptom severity).* M = Male, F = Female, NB = Non-binary, T = Transgender. Other = “none of the above”. \*Significant differences between responder status groups; <sup>+</sup> Significant differences between mechanism groups.

### Unmedicated Control Comparisons

To isolate antidepressant-associated effects beyond current symptom severity, R and NR were each matched to unmedicated participants on current depression severity using propensity score matching (details below). This yielded matched unmedicated control groups for R (UMC-R; N = 40) and NR (UMC-NR; N = 17). Participant demographics for the matched unmedicated control groups are provided in Supplementary Materials (Table S1).

### Questionnaires

We administered a battery of questionnaires to assess depression severity, comorbid psychiatric symptoms, medical and treatment history, and other relevant factors (see Table 1 and Supplemental Methods for full list).

### Emotional mnemonic discrimination task

Participants completed an established emotional MDT during fMRI scanning (21,25) (Fig. S1). During encoding, participants viewed negative, neutral, and positive images and provided valence ratings (negative, neutral, or positive). After a brief delay, memory was tested using repeated images (targets), similar images (lures), and novel images (foils). Participants indicated whether each image was “Old” or “New/Different.” Analyses focused on correct rejections (CRs) of lure stimuli, which are thought to reflect signals consistent with hippocampal pattern separation (25). Analyses of the lure false alarms (FAs; falsely categorizing a lure as ‘Old’) are included in the Supplementary Materials (Supplementary Results 3, 6, and 10). Behavioral performance measures are reported in Table 1, with additional analyses provided in the Supplementary Materials (Table S1).

### High-resolution structural and functional neuroimaging data acquisition

We collected high resolution structural and functional MRI scans using a Siemens Prisma 3T scanner (32-ch head coil). We collected whole-brain high resolution T1-weighted structural scans (0.8mm isotropic) [Orientation = Sagittal; Slices per slab = 208; FoV = 256mm; Slice thickness = 0.8mm; TR = 2400ms; TE = 2.24ms; Flip angle = 8 deg; PAT mode = GRAPPA; Phase encoding direction = A > P; Multi-slice mode = single shot; Series = Interleaved]. To apply distortion correction, two spin-echo EPI images, one with anterior-to-posterior (AP) phase encoding and one with posterior-to-anterior (PA) phase encoding, were acquired before each BOLD scan. Whole-brain functional MRI BOLD scans (1.5mm isotropic) were acquired [Orientation = Parallel to the hippocampus; Slices per slab = 88; FoV = 212mm; Slice thickness = 1.54mm; TR = 2400ms; TE = 35.60ms; Multiband accel. factor = 4; Flip angle = 45 deg; PAT mode = GRAPPA; Phase encoding direction = A > P; Multi-slice mode = interleaved; Series = Interleaved] during each phase of the emotional MDT. Neuromelanin-sensitive locus coeruleus scans and resting-state MRI scans were also collected, although they were not utilized for the current study.

### Neuroimaging data analysis and regions of interest

Imaging preprocessing and analysis was conducted using AFNI version 24.0.17 software and analytical pipelines adapted from previous work (25,40,41). Functional images were first corrected for distortion using the unWarpEPI.py function. Slice-timing and motion correction were performed using the Fourier interpolation method. Significant motion events, or time points in which motion exceeded 0.5 mm in translation across any axis, relative to prior acquisition ±1 time point, were censored from subsequent analyses. For similar reasons, de-spiking was applied to images using the default parameters from the 3dDespike function to further correct for potential physiological noise. Next, functional masks were created from corrected images then aligned to individual skull-stripped structural scans. Alignment was visually inspected for each scan, and manual adjustments were made when necessary. To enhance our signal-to-noise ratio, we performed smoothing with a Gaussian kernel of 2.0 mm full width at half maximum (FWHM).

Our primary regions of interest (ROIs) were the hippocampal DG/CA3 subfields and the BLA based on prior work (25), the subfields’ role in the emotional modulation of memory (25), as well as their susceptibility to antidepressant effects (42). We further stratified hippocampal ROIs by anterior and posterior axis. We included CA1 and the central nucleus of the amygdala (CEA) as control regions (Fig. 1). ROIs were defined using previously established anatomical segmentation protocols (43). Hippocampal subfield masks were based on prior work from Yassa et al. (2010) (44–46) defined by the atlas of Duvernoy. In this protocol, hippocampal subfields are delineated on eight coronal slices spanning the anterior-posterior axis of the hippocampus. Representative slices are first identified based on correspondence with atlas landmarks, and segmentation is then extended slice-by-slice in both directions to ensure smooth transitions across slices. Amygdala subnuclei were defined according to Entis et al. (2012) (47), using anatomical landmarks including the hippocampal uncus, temporal horn of the lateral ventricle, hippocampus, and adjacent white matter to delineate the basolateral and central nuclei.

**Figure 1.**
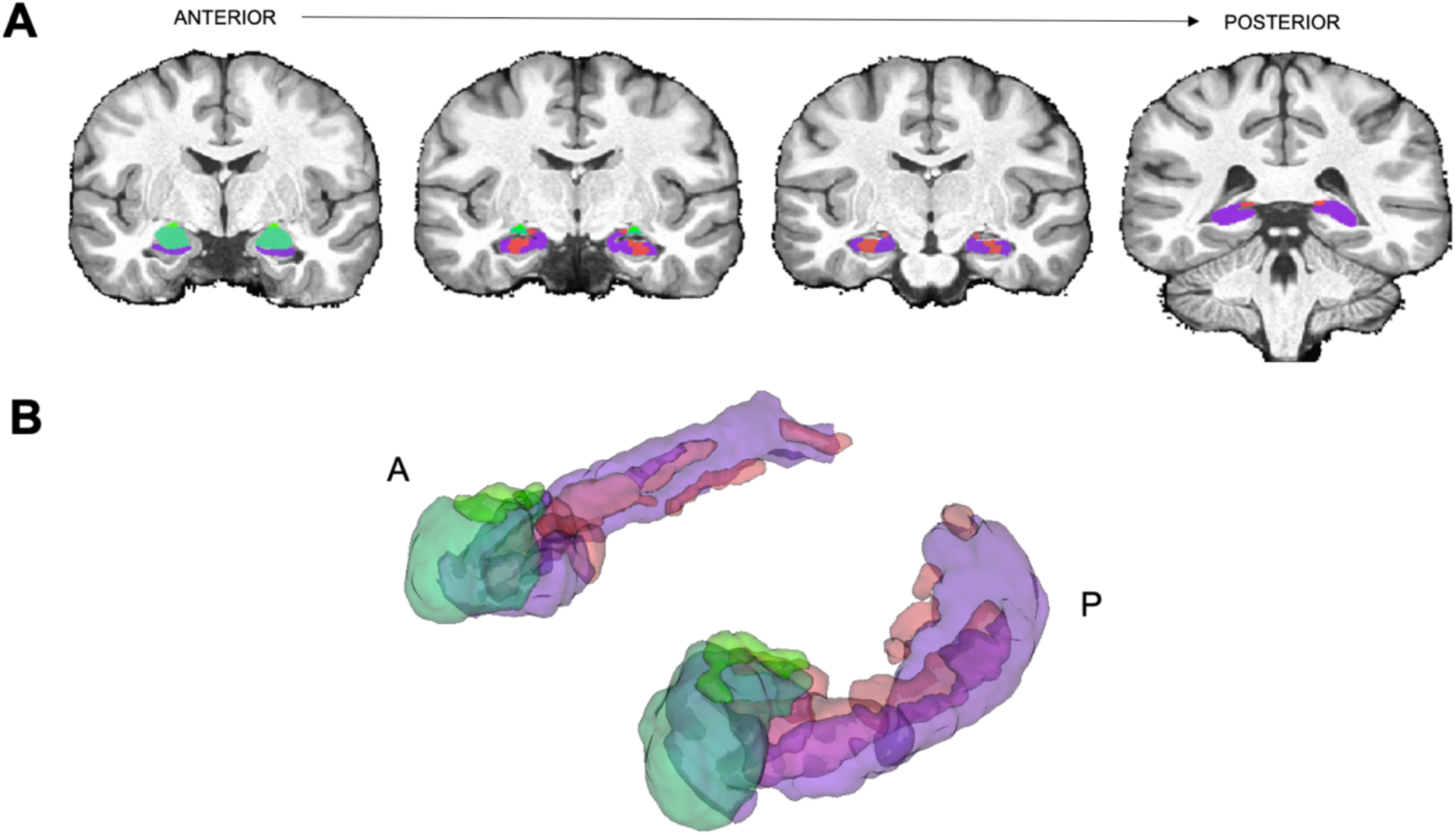
Hippocampal and amygdala subfield regions of interest. A) ROI segmentations of the amygdala (BLA, CEA) and hippocampal subfields (DG/CA3, CA1) generated on a custom group-level template. B) A schematic of a 3D rendering of our high-resolution anatomical template including the dentate gyrus (DG)/CA3 (red), CA1 (purple), basolateral amygdala (BLA; teal), and central nucleus of the amygdala (CEA, light green).

Masks were manually delineated on the high-resolution anatomical template (.065 mm isotropic) and visually inspected to ensure anatomical accuracy. We adopted this template-based approach to maximize consistency across participants and minimize variability that can arise when manually segmenting small MTL structures in each individual dataset. Manual segmentation in native space is highly labor-intensive and can introduce subject-specific variability in boundary placement, particularly in regions where tissue contrast is limited. By defining ROIs once on a high-resolution template and applying the same masks to all participants, we ensured that identical anatomical criteria were used across the entire sample.

Structural scans were registered to a high-resolution template space (.065 mm) using Advanced Normalization Tools (ANTS) (48) before performing deconvolution procedures. Functional voxel masks were combined with the anatomical ROI masks to extract activity only from voxels falling within each ROI. All registrations were visually inspected and manually adjusted when necessary to ensure accurate alignment of hippocampal and amygdala subregions. To model task-related activity, condition-specific regressors (“behavioral vectors”) were generated from stimulus onset times for each trial type, categorized by emotional valence (negative, neutral, positive) and memory outcome (e.g., hits, lure correct rejections [lure CRs], lure false alarms [lure FAs], and correct rejections to novel foils). These regressors were entered into a voxel-wise general linear model (GLM) implemented in AFNI using 3dDeconvolve. Stimulus time courses were modeled using multiple linear regression to estimate the hemodynamic response for each condition relative to a baseline of novel foil CRs. This approach is commonly used and is effective for isolating neural activity associated with successful recollection and pattern separation (25,40,49). Given the visual similarity between lure trials and novel foils, differences in signal are attributable primarily to memory-related processing rather than low-level perceptual differences. Using an active baseline condition is also generally preferred over passive baselines such as fixation or rest (50). For each trial type, beta coefficients were summed across the expected hemodynamic response window (3-12 seconds post-stimulus) to generate a single estimate of neural activity for each ROI. Activity during lure CRs and lure FAs was averaged across bilateral ROIs for all subsequent analyses. We focused our primary analyses on lure CRs because these trial types most closely index hippocampal pattern separation, the central mechanism motivating our hypotheses. In contrast, conventional recognition memory contrasts such as Hits versus Misses reflect broader episodic memory processes and are less specific to DG/CA3-dependent discrimination of highly similar stimuli.

### Statistical Analyses

Analyses were conducted using R studio 4.5.0 and JASP 0.19.1. We conducted mixed ANOVAs, *t*-tests (two-tailed), regressions, Pearson correlations, and Fisher’s r-to-z transformations when relevant. Post hoc comparisons were conducted using two-tailed independent- and paired-samples t-tests, as appropriate. When omnibus ANOVA models included factors with more than two levels, Scheffé’s tests were used to examine significant effects. For all other planned post hoc comparisons, p-values were adjusted for multiple comparisons using the Holm-Bonferroni sequential correction procedure. All statistical tests, ANOVAs and correlations, used the General Linear Model. Sphericity violations in repeated-measures tests were corrected using the Greenhouse-Geisser correction. Welch’s correction was applied for unequal variances, when relevant. Statistical values were controlled for Type 1 errors and considered significant at an alpha level of 0.05. Any result referred to as marginal was considered to be marginal an alpha level of 0.08. Unmedicated controls were matched separately to responders and non-responders using propensity score matching implemented in the R package *MatchIt*. Propensity scores were estimated using a logistic regression model with current depressive symptom severity (BDI-II total score) as the sole matching covariate. Nearest-neighbor matching was performed with a caliper of 0.25 and a 1:1 matching ratio without replacement. This analysis was conducted separately for responders and non-responders, with no overlap in unmedicated controls across the two matched samples (51). We selected current depressive symptom severity as the only matching variable because it represented the primary potential confound of interest. This approach was designed to test whether neural differences associated with antidepressant use persisted after accounting for current symptom burden, rather than to create groups balanced across all demographic and clinical characteristics. Consequently, groups were not expected to be equivalent on other variables such as age, education, or anxiety symptoms. These variables were examined and are reported in the Supplementary Materials for transparency.

To determine how responder status and antidepressant mechanism were associated with hippocampal and amygdala activity, we conducted a series of mixed ANOVAs. Primary inferences were based on models that included both responder status and antidepressant mechanism as between-subjects factors and emotion and region as within-subject factors. Hippocampal (DG/CA3, CA1) and amygdala (BLA, CEA) subregions were modeled together to enable within-subject comparisons and to assess the specificity of effects in our *a priori* regions of interest (DG/CA3 and BLA) relative to comparison regions (CA1 and CEA). We conducted the above models in anterior and posterior hippocampal regions. Additionally, given that the responder-by-mechanism interaction included unequal group sizes, particularly the small number of non-responders taking multi-action antidepressants, these effects were interpreted cautiously. To aid interpretation of significant interaction effects, we conducted exploratory follow-up models examining responder status alone (R, n = 47; NR, n = 22) and antidepressant mechanism alone (Single-action, n = 37; Multi-action, n = 32). These models provided greater statistical power to detect the independent effects of each factor on hippocampal and amygdala activity during negative, neutral, and positive lure CRs.

We applied the same analytic framework to examine task-related hippocampal-amygdala coactivation (i.e., correlations between beta weights across regions) during lure CRs. Finally, to isolate antidepressant-associated effects beyond current symptom severity, we compared medicated participants with unmedicated controls matched on current depression severity (R: n = 40 vs. UMC-R: n = 40; NR: n = 17 vs. UMC-NR: n = 17) using the same set of analyses. The data generated in the current study are available in a GitHub repository (https://github.com/lealmemorylab/Antidepressant-Imaging).

## Results

### No differences in participant characteristics by mechanism, with expected symptom-measure differences in responder status

First, we examined demographic and questionnaire data across responder groups (R, NR) and mechanism groups (single-action, multi-action). Across responder groups, there were significant group differences across R and NR on BDI-II (current) [*t*(67) = 8.35, *p* < .001, *d* = 2.16], BAI (current) [*t*(67) = 2.85, *p* = .01, *d* = 0.74], current levels of stress [*t*(67) = 5.67, *p* < .001, *d* = 1.47], and past month levels of stress [*t*(67) = 4.24, *p* < .001, *d* = 1.10], where NRs reported greater depressive symptoms, anxiety symptoms, and stress levels compared to R, as expected. Importantly, there were no differences in retrospective BDI-II across groups, suggesting R and NR started at similar depression levels [*t*(67) = 1.29, *p* = .203, *d* = 0.33]. Across mechanism groups, there was a significant difference between single- and multi-action groups in antidepressant switching frequency [*t*(44) = 2.57, *p* = .014, *d* = −0.64], in which the multi-action mechanism group reported more frequent switching of antidepressants than the single-action mechanism group. Importantly, there were no differences in retrospective BDI-II across groups, suggesting those taking single- or multi-action antidepressants started at similar depression levels [*t*(67) = 0.17, *p* = 869, *d* = −0.04]. Task accuracy did not differ by responder status or mechanism; however, across all analyses, there was a main effect of emotion such that discrimination was higher for neutral relative to emotional lures, in line with prior work (21). Detailed group comparisons and analyses are reported in Supplemental Results 1. Group comparisons to unmedicated controls are reported in Supplemental Results 1 and in Table S1.

### Responder status and mechanism interact to differentially impact anterior hippocampal subfield activity

First, we conducted a mixed ANOVA on lure CR activity with emotion (Negative, Neutral, Positive) and region (DG/CA3, CA1) as within-subject factors and both responder status (R, NR) and mechanism (single-action, multi-action) as between-subjects factors in the anterior hippocampus. There was a significant three-way interaction between region x mechanism x responder status [*F*(1, 65) = 4.21, *p* = .044, 17^2^ = 0.004], such that NR taking single-action antidepressants showed greater activity in DG/CA3 relative to CA1, whereas NR taking multi-action antidepressants showed greater activity in CA1 relative to DG/CA3 [*F*(1, 65) = 4.75, *p* = .033, 17^2^ = 0.07] (Fig. 2A-B, S2 for interaction visualization), with no such differences in R [*p* > .05]. However, this interaction should be interpreted with caution given the small sample size of those taking multi-action antidepressants who were non-responders. To aid interpretation and increase power within responder status and mechanism groups independently, we conducted exploratory responder-only and mechanism-only models. Both models revealed a main effect of region (DG/CA3 > CA1), but no interactions involving responder status or mechanism (Supplemental Results 2). Follow-up analyses examining posterior hippocampal subfields and hippocampal axis specificity (anterior vs. posterior) provided little evidence for comparable effects outside the anterior hippocampus (Supplemental Results 2; Fig. S3).

**Figure 2.**
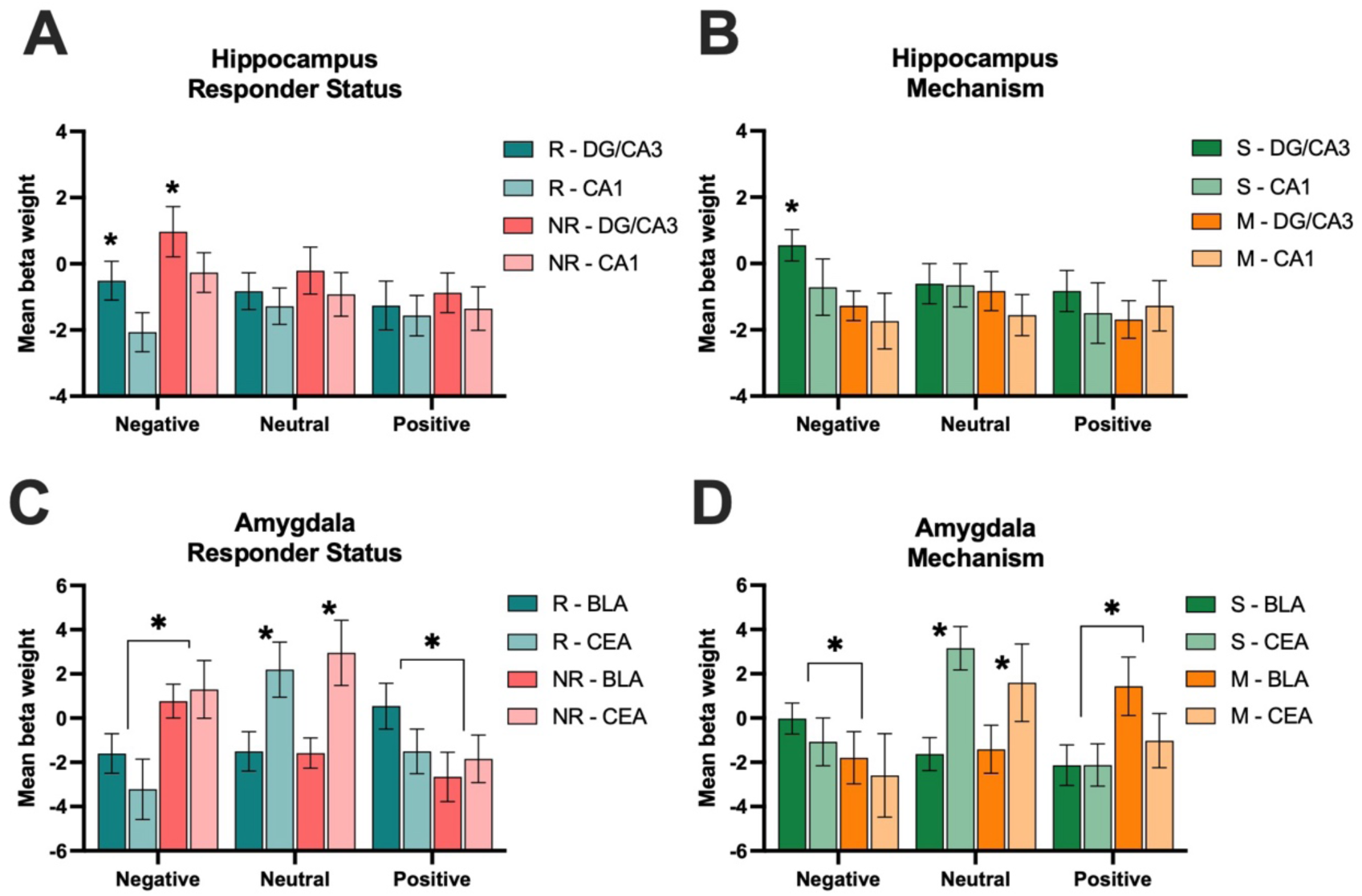
Anterior hippocampal (DG/CA3, CA1) and amygdala subnuclei (BLA, CEA) activity based on responder status and mechanism. A) Mean anterior hippocampal activity beta weights during lure CRs across responder status and region. B) Mean anterior hippocampal activity beta weights during lure CRs across mechanism and region (*p < .05 and represents main effect of region, DG/CA3 > CA1). C) Mean amygdala activity beta weights during lure CRs across responder status. D) Mean amygdala activity beta weights during lure CRs across mechanism (*p < .05 and represents an emotion by mechanism interaction and an emotion by region interaction).

### Responder status and mechanism show independent interactions with emotion in the amygdala

Next, we conducted a mixed ANOVA on lure CR activity with emotion (Negative, Neutral, Positive) and region (BLA, CEA) as within-subject factors and both responder status (R, NR) and mechanism (single-action, multi-action) as between-subjects factors. There was a significant emotion x responder status interaction [*F*(2, 130) = 4.01, *p* = .020, 17^2^ = 0.019], where R showed less amygdala activity during negative lure CRs compared to NR but greater amygdala activity during positive lure CRs [*F*(1,65) = 6.48, *p* = .013, 17^2^= 0.091] (Fig. 2C-D, S6A for interaction visualization). There was also a significant emotion x region interaction [*F*(2, 130) = 9.72, *p* < .001, 17^2^ = 0.016], where there was greater activity during neutral lure CRs in the CEA compared to the BLA [*F*(1,65) = 22.05, *p* < .001, 17^2^ = 0.253] (Fig. 2C-D, S6B for interaction visualization). Exploratory responder-only and mechanism-only models largely recapitulated effects observed in the primary model. An additional emotion x mechanism interaction emerged in the mechanism-only model, whereby individuals taking single-action antidepressants showed greater amygdala activity during negative lure CRs and lower activity during positive lure CRs relative to those taking multi-action antidepressants (see Supplemental Results 4 for full statistical models).

### Greater hippocampal-amygdala coactivation during negative lure discrimination in responders and those taking multi-action antidepressants

Next, we tested whether responder status and mechanism influenced functional coactivation of hippocampal subregions (DG/CA3, CA1) and amygdala subnuclei (BLA, CEA). Correlation matrices were generated using pairwise r values across ROIs for each emotion (Negative, Neutral, Positive) and compared 1) R and NR, and 2) single-action and multi-action mechanism groups hippocampal-amygdala functional coactivation. We were not powered to examine groups stratified at the level of responder status and mechanism given the small number of non-responders taking multi-action antidepressants. We also created a contrast correlation matrix representing the difference in pairwise r values between groups. All correlations were normalized using Fisher’s r-to-z transformation and compared using a z-difference test. We corrected for multiple comparisons using Bonferroni for each correlogram (adjusted *p* = .0083 threshold). Analyses revealed that R exhibited greater anterior DG/CA3-BLA coactivation during negative lure CRs (*r* = 0.588, *p* < .001) compared to NR (*r* = −0.051, *p* = .822) [*z* = 2.64, *p* = .008] (Fig. 3A-C). There were no other significant group differences in functional coactivation for any other ROI pairs or within neutral and positive lure CRs (Fig. 3D-I). Applying the same analyses to posterior hippocampal regions revealed no significant differences across responder status (Supplemental Results 5, Fig. S7). Similarly, no significant effects were observed when analyses were conducted across the whole hippocampus without stratifying by anterior-posterior axis.

**Figure 3.**
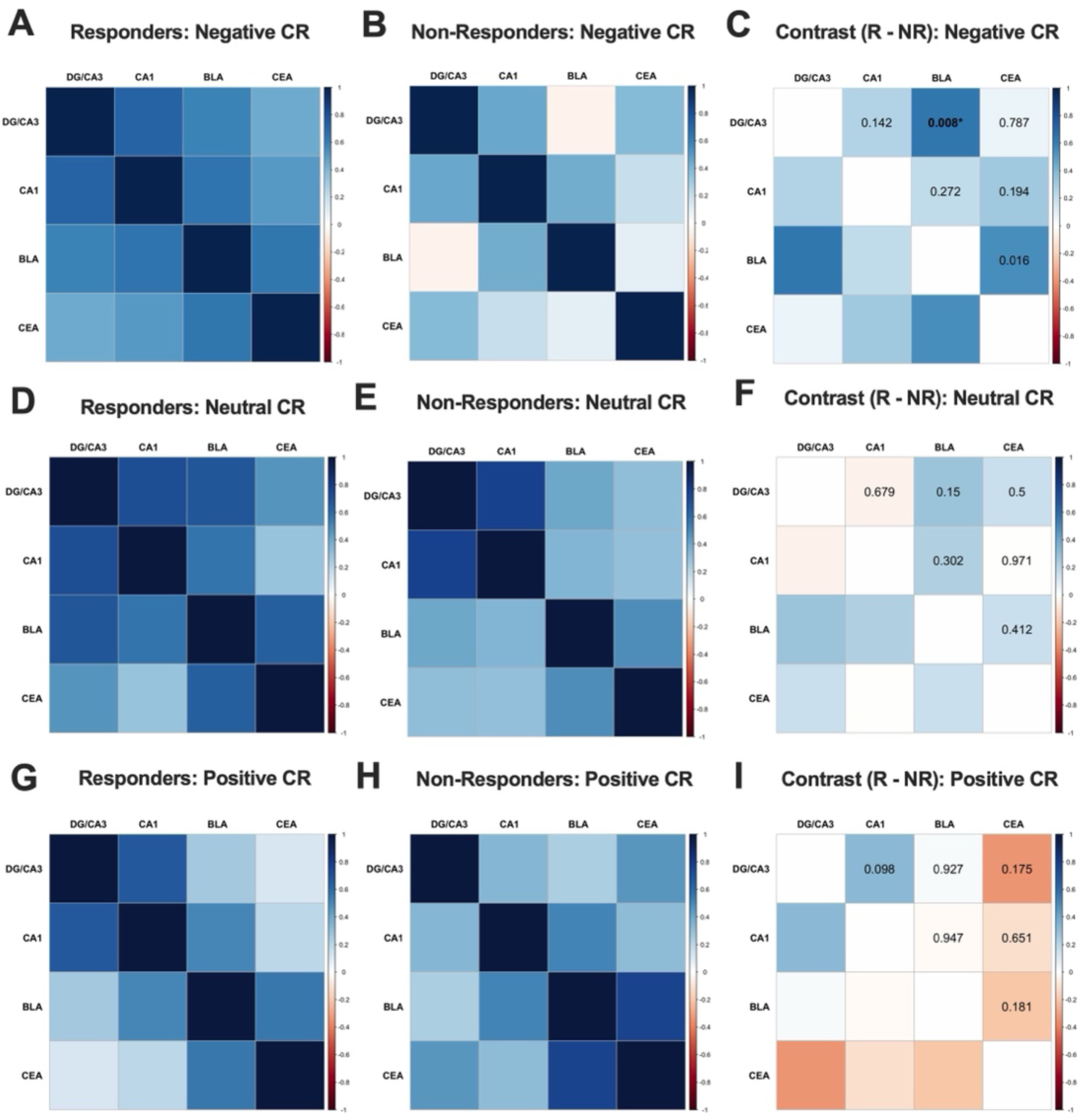
Hippocampal-amygdala functional coactivation during lure CRs in responders and non-responders. A) Negative lure CR correlation matrices for pairwise r values across anterior hippocampal (DG/CA3, CA1) and amygdala (BLA, CEA) regions of interest (ROIs) in responders (R) and B) non-responders (NR); C) Negative lure CR contrast correlation matrix representing the difference in pairwise r values between R and NR. D) Neutral lure CR correlation matrices for pairwise r values across ROIs in R and E) NR groups; F) Neutral lure CR contrast correlation matrix representing the difference in pairwise r values between R and NR. G) Positive lure CRs correlation matrices for pairwise r values across ROIs in R and H) NR; I) Positive lure CRs contrast correlation matrix representing the difference in pairwise r values between R and NR. The contrast correlation matrices were normalized using Fisher’s r-to-z transformation and compared using a z-difference test (*p < .0083; corrected for multiple comparisons).

Next, we examined whether hippocampal-amygdala functional coactivation differed as a function of antidepressant mechanism. Analyses revealed that the multi-action mechanism group exhibited greater DG/CA3-BLA (*r* = 0.500, *p* < .001), BLA-CA1 (*r* = 0.594, *p* < .001), and BLA-CEA (*r* = 0.800, *p* < .001) functional coactivation during negative lure CRs compared to the single-action mechanism group (DG/CA3-BLA: *r* = 0.295, *p* = 0.853; BLA-CA1: *r* = 0.393, *p* = 0.116; BLA-CEA: *r* = 0.282, *p* = 0.167) [DG/CA3-BLA: *z* = −3.68, *p* < .001; BLA-CA1: *z* = −3.16, *p* = .002; BLA-CEA: *z* = −2.72, *p* = .007] (Fig. 4A-C). There were no other significant differences in functional coactivation for any other ROI pairs or within neutral and positive emotion lure CRs (Fig. 4D-I). We were underpowered to stratify R and NR by mechanism type; however, we included these exploratory analyses and results in the Supplement (Fig. S8-9).

**Figure 4.**
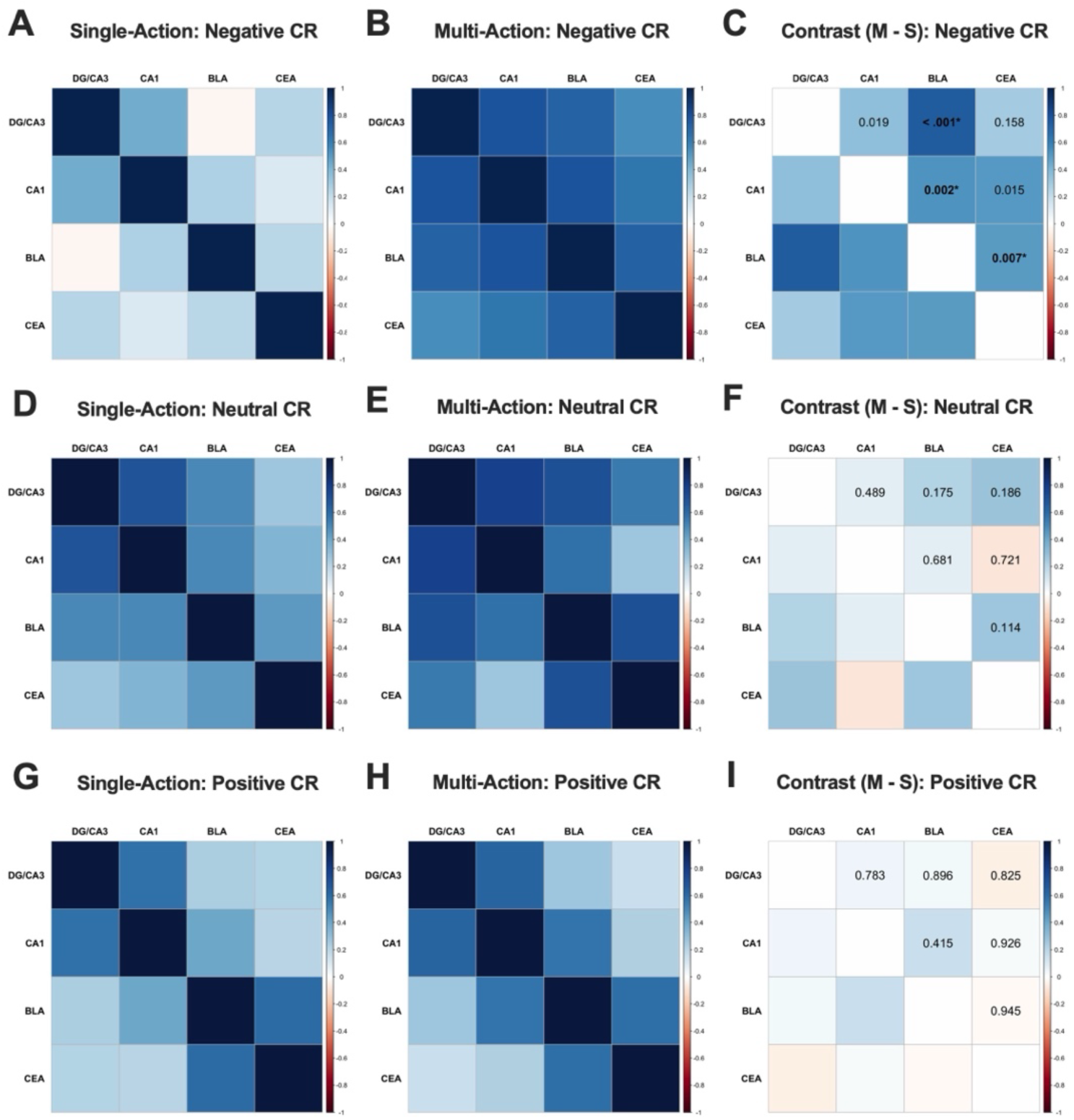
Hippocampal-amygdala functional coactivation during lure CRs in single-action and multi-action mechanism groups. A) Negative lure CR correlation matrices for pairwise r values across anterior hippocampal (DG/CA3, CA1) and amygdala (BLA, CEA) regions of interest (ROIs) in single-action and B) multi-action groups; C) Negative lure CR contrast correlation matrix representing the difference in pairwise r values between single-action and multi-action groups. D) Neutral lure CR correlation matrices for pairwise r values across ROIs in single-action and E) multi-action groups; F) Neutral lure CR contrast correlation matrix representing the difference in pairwise r values between single-action and multi-action groups. G) Positive lure CRs correlation matrices for pairwise r values across ROIs in single-action and H) multi-action groups; I) Positive lure CRs contrast correlation matrix representing the difference in pairwise r values between single-action and multi-action groups. The contrast correlation matrices were normalized using Fisher’s r-to-z transformation and compared using a z-difference test (*p < .0083; Bonferroni corrected for multiple comparisons).

Applying the same analyses to posterior hippocampal regions revealed no significant differences across mechanism (Fig. S10). Follow-up analyses examining whole-hippocampus connectivity largely converged with the primary findings, with the multi-action antidepressant group showing greater negative emotional DG/CA3-centered functional coactivation than the single-action group (see Supplemental Results 5).

### Antidepressant-associated differences beyond current depression severity across responders and unmedicated matched controls in the amygdala

To determine whether the observed effects reflected antidepressant-associated differences beyond current depression severity, we compared R to UMC-R matched on current depression severity. There was a main effect of region [*F*(1, 78) = 11.45, *p* = .001, 17^2^ = 0.006] (Fig. 5A), where there was greater DG/CA3 activity during lure CRs relative to CA1, but no significant main effects of group nor interactions with group regardless of hippocampal axis (Supplemental Results 7), suggesting that hippocampal activity in R was not significantly different than unmedicated individuals matched on depression severity. We also performed exploratory analyses stratifying the R by single-action and multi-action and compared to the UMC-R, but found no significant effects or interactions with group (Supplemental Results 7).

**Figure 5.**
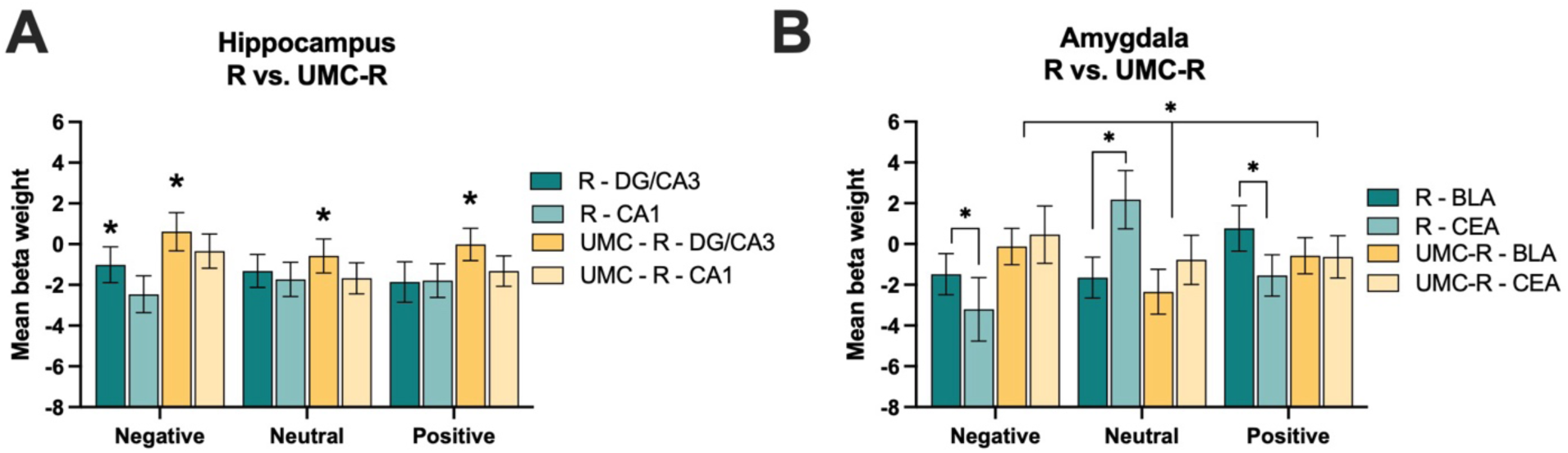
Anterior hippocampal and amygdala activity in responders and unmedicated controls matched on depression symptom severity. A) Mean activity beta weights in hippocampal subregions during lure CRs across responders (R) and unmedicated matched controls (UMC-R) groups. B) Mean activity beta weights in amygdala subregions during lure CRs across R and UMC-R groups (*p < .05).

In the amygdala, we conducted a mixed ANOVA during lure CR activity with emotion (Negative, Neutral, Positive) and region (BLA, CEA) as within-subject factors and group (R, UMC-R) as a between-subjects factor. There was a significant three-way interaction between emotion x region x group [*F*(2, 156) = 3.58, *p* = .030, 17^2^ = 0.005], where R show greater BLA compared to CEA activity during emotional lure CRs but the opposite pattern during neutral lure CRs and in contrast to UMC-R who showed greater amygdala activity during emotional lure CRs compared to neutral regardless of amygdala subnuclei [*F*(1,78) = 7.62, *p* = .007, 17^2^ = .089] (Fig. 5B). There was also a marginally significant emotion x group interaction [*F*(2,156) = 3.03, *p* = .051, 17^2^ = 0.001], where R had less activity during negative lure CRs and greater activity during neutral lure CRs compared to UMC-R [*F*(1,78) = 4.67, *p* = .034, 17^2^ = 0.056]. There was also a significant interaction between emotion and region [*F*(2, 156) = 9.10, *p* < .001, 17^2^ = 0.013], where there was greater CEA activity relative to BLA activity during neutral lure CRs compared to emotional lure CRs [*F*(1,78) = 18.97, *p* < .001, 17^2^ = 0.196]. This suggests that antidepressants showed a drug-associated difference beyond current severity on amygdala activity in R (e.g. emotion, region, group interaction). We also performed exploratory analyses stratifying the R by single-action and multi-action and compared to the UMC-R and found marginal interactions with group (Supplemental Results 7).

When we examined whether hippocampal-amygdala functional coactivation differed between R and UMC-R, there were no significant group differences in hippocampal-amygdala functional coactivation regardless of hippocampus axis (Fig. S14). Finally, we compared NR to UMC-NR matched on current depression severity (e.g. untreated depression) and found no group differences in hippocampal and amygdala activity (Supplemental Results 9, Fig. S15) nor coactivation (Fig. S16).

## Discussion

Antidepressants remain central to depression treatment, yet their effects on the cognitive and neural systems that are disrupted in mood disorders remain poorly understood in humans. Much of what is known comes from animal models demonstrating that monoaminergic modulation, neuroplasticity, and hippocampal neurogenesis shape emotional memory and stress responses. However, little human work has clarified how antidepressants alter computations within hippocampal-amygdala circuits that contribute to emotional memory and depressive symptomatology, particularly those governing hippocampal pattern separation and emotional mnemonic discrimination. By applying a high-resolution, subfield-level approach during an emotional MDT, the present study identifies distinct neural signatures associated with antidepressant mechanism, treatment response, and their interaction, helping bridge systems-level neurobiology and individualized treatment strategies.

### Antidepressant efficacy and mechanism shape separable hippocampal computation profiles

Across analyses within the anterior hippocampus, DG/CA3 activity exceeded CA1 activity during lure discrimination, consistent with the known role of DG/CA3 in pattern separation (25). Critically, subfield recruitment varied by medication mechanism and treatment response. NR taking single-action antidepressants showed DG/CA3 hyperactivity, whereas NR taking multi-action medications showed the opposite pattern and a relative shift toward CA1 engagement. This crossover interaction suggests that non-response reflects distinct disruptions in anterior hippocampal computations involved in emotional memory processing linked to pharmacological targets. Ineffective serotonergic modulation may destabilize DG/CA3 excitatory-inhibitory dynamics (52,53), whereas multi-action antidepressants influencing norepinephrine and dopamine systems that project more directly to CA1 (54), may bias processing towards CA1. This relative shift toward CA1 engagement may reflect hippocampal processing that favors generalization over discrimination. This shift could lead to less distinct encoding of similar emotional experiences, potentially promoting overgeneralized negative memory representations in non-responders. Hippocampal subfield balance may be a potential mechanistic target for interventions aimed at improving emotional memory biases in depression.

Notably, these effects were localized to the anterior hippocampus, with no comparable effects in posterior hippocampus. Given the stronger connectivity of the anterior hippocampus with the amygdala and its established role in emotional memory processing, this pattern suggests that antidepressant-related differences in hippocampal subfield function during emotional mnemonic discrimination may be preferentially expressed within anterior hippocampal circuitry. Consistent with this interpretation, prior human work indicates that antidepressant treatment is associated with increased neural progenitor cell number in the anterior DG (55), although the extent to which this process is necessary for therapeutic effects remains unclear. Together, these findings suggest that antidepressant efficacy may be linked to restoration of DG-dependent computations supporting emotional pattern separation and hippocampal-amygdala circuit function.

In contrast, R showed a canonical DG/CA3 > CA1 profile, suggesting successful treatment normalizes hippocampal computations during lure discrimination. This pattern aligns with neuroplasticity models positing that effective antidepressants enhance DG neurogenesis, restore CA3 dendritic integrity, and rebalance hippocampal-amygdala interactions (3). Although the responder-by-mechanism interaction was statistically significant, only five non-responders were taking multi-action antidepressants. As a result, this finding should be considered exploratory until replicated in larger samples. To aid interpretation of this underpowered interaction, we also examined responder-only and mechanism-only models, which were better powered to detect the independent effects of treatment response and antidepressant mechanism. Future work examining specific antidepressant types will clarify how distinct neurotransmitter profiles shape subfield dynamics. Together, these findings identify hippocampal subfield activity as a mechanistically specific biomarker of treatment response and highlight the importance of antidepressant mechanism in determining neural outcomes.

### Emotion-guided amygdala signaling dissociates responders from non-responders

Amygdala activity showed effects of both treatment response and antidepressant mechanism that were specific to emotional valence. Importantly, the emotion by responder status interaction was significant in both the full model and the responder-only follow-up model, indicating that this effect was robust to the inclusion of antidepressant mechanism. Responders exhibited reduced amygdala activity during negative lure discrimination but increased activity during positive lure discrimination, consistent with normalization of valence-sensitive emotional memory and reduced negative memory bias following effective treatment (56–58). Notably, responder profiles also differed from unmedicated, symptom-matched controls, indicating that these effects likely reflect antidepressant-associated neural changes beyond current depression severity.

In contrast, the emotion by mechanism interaction was significant only in the mechanism-only follow-up model and did not remain significant in the full model that simultaneously accounted for responder status. This pattern suggests that mechanism-related differences were smaller in magnitude and more sensitive to reduced statistical power associated with unequal subgroup sizes. Accordingly, these findings should be considered exploratory. Within this context, individuals taking single-action antidepressants showed greater amygdala responses to negative lures and lower responses to positive lures compared to those taking multi-action medications, consistent with differential serotonergic and catecholaminergic modulation of amygdala circuits. Together, these findings suggest that antidepressants do not uniformly dampen amygdala activity, but rather may reshape emotion-specific computations in ways that depend on treatment efficacy and, more tentatively, pharmacological mechanism.

### Coactivation markers reveal a circuit-level signature of effective treatment

Analyses identified heightened anterior DG/CA3-BLA functional coactivation during negative lure discrimination in responders and individuals taking multi-action antidepressants. Given the strong anatomical and functional connectivity between anterior hippocampus and the amygdala (59), this pattern suggests that effective antidepressant treatment may preferentially restore emotional memory circuitry involved in emotional pattern separation. This effect was absent in NR and symptom-matched unmedicated controls, indicating specificity to effective treatment. Although DG/CA3-BLA showed the strongest coactivation effect and was the only hippocampal-amygdala connection to survive correction for multiple comparisons in responder analyses, direct comparisons across circuits did not reveal statistically significant differences in effect magnitude relative to other hippocampal subfield or amygdala subnuclei pairings. Accordingly, these findings suggest relatively greater involvement of the DG/CA3-BLA pathway, rather than exclusive or selective engagement of this circuit. These findings suggest that effective antidepressant treatment may restore hippocampal-amygdala coactivation disrupted in depression (60,61).

Mechanism-related differences in coactivation suggested that individuals taking multi-action antidepressants showed broader hippocampal-amygdala coactivation (e.g. DG/CA3-BLA, BLA-CA1, and BLA-CEA) during negative lure discrimination compared to those taking single-action antidepressants. If replicated, this pattern would be consistent with models proposing that catecholaminergic modulation enhances integration of emotional salience into memory (62,63). Notably, effects were specific to negative stimuli, consistent with preferential engagement of hippocampal-amygdala circuits during emotionally salient information, with effects being more pronounced for negative relative to positive emotional stimuli (64). Greater coactivation did not simply reflect increased activity, as R showed stronger hippocampal-amygdala coactivation despite lower regional activation. Together, these findings identify hippocampal-amygdala coactivation during negative emotional discrimination as a candidate circuit-level biomarker of treatment response.

### Disentangling drug effects from symptom severity

Antidepressant response is highly variable, making it critical to identify neural signatures that distinguish R from NR. Comparing R to unmedicated individuals matched on depression severity allowed us to isolate treatment-related neural effects beyond current symptoms. R showed no differences in hippocampal activity or hippocampal-amygdala coactivation relative to matched unmedicated controls, suggesting normalization of hippocampal function with effective treatment.

In contrast, amygdala activity differed, in which R exhibited greater BLA relative to CEA activity during emotional lure discrimination and greater CEA relative to BLA activity during neutral lure discrimination, whereas unmedicated controls showed elevated amygdala activity during emotional lures regardless of subnucleus. This pattern indicates that antidepressants selectively modulate amygdala function, supporting more balanced emotional memory processing beyond symptom severity.

Mechanism-related differences further suggested that R taking multi-action antidepressants showed a more adaptive profile (e.g. reduced negative and enhanced positive BLA engagement) compared to single-action R and unmedicated controls. This pattern is consistent with broader neurotransmitter modulation shifting amygdala processing away from a negativity bias. While prior research has found that single-action antidepressants such as SSRIs can reduce amygdala hyperactivity in response to negative stimuli, there is less consensus on whether this effect extends to other mechanisms of action (65). However, these exploratory effects were modest, and larger samples are needed to clarify mechanism-specific influences on amygdala subnuclei.

### Translational implications: Toward mechanistically informed precision psychiatry

These findings have several implications for personalized antidepressant treatment. First, if validated prospectively, mechanism-specific biomarkers within anterior hippocampal circuitry, such as DG/CA3 versus CA1 activity profiles, may help identify patients mismatched to their current antidepressant mechanism. Second, anterior DG/CA3-BLA coactivation during negative memory may serve as a sensitive biomarker for monitoring treatment-related neurobiological change. Third, identifying DG/CA3 hyperactivity as a marker of ineffective treatment suggests that interventions targeting DG function (e.g. exercise, neuromodulatory interventions, drug-induced plasticity mechanisms, etc.) may improve outcomes. Finally, because effects were strongest for negative stimuli, emotion-focused memory tasks may may provide sensitive cognitive markers for assessing treatment efficacy.

### Limitations

Several limitations warrant consideration. First, although powered to compare single- and multi-action antidepressants, heterogeneity within the multi-action group, particularly across drug type and responder status, limited our ability to isolate effects of NE- or DA-selective agents. Larger, stratified samples will be needed to clarify neurotransmitter-specific mechanisms. Second, the cross-sectional design also limits causal inference, as observed neural differences may reflect pre-existing vulnerabilities rather than treatment effects. We relied on retrospective self-reports of pre-treatment symptoms, and while generally reliable (35,36,66), remain vulnerable to recall and state-dependent bias. Prospective longitudinal designs with pre-treatment baselines would allow more precise assessment of biomarker sensitivity and reduce concerns such as regression to the mean (67). Third, responder classification also varies depending on the scale used and distinctions between response and remission (68). Because our focus was perceived treatment efficacy, future studies should incorporate standardized clinical criteria to refine responder definitions. Fourth, comorbidity further complicates interpretation. Over half of participants reported co-occurring disorders (e.g., anxiety, OCD, ADHD), which enhances generalizability but may introduce variability, as comorbid symptoms can influence treatment response and emotional memory circuits (30,69–72). Future work should examine how comorbidity and depression subtypes shape antidepressant-related neural effects. Finally, although we focused on hippocampal and amygdala subregions, depression involves broader brain networks, including prefrontal regions implicated in emotion regulation and hippocampal coactivation (23,73–75). Future studies incorporating whole-brain and network-level approaches will be important for contextualizing these regionally specific findings within distributed neural systems involved in affective and cognitive processing.

## Conclusions

Antidepressants are the primary pharmacological treatment for depression, yet their circuit-level mechanisms in humans remain poorly understood, particularly regarding pattern separation. These findings identify circuit-level biomarkers that may support more personalized, mechanistically informed treatment. This approach advances precision psychiatry by enabling antidepressant selection and monitoring based on measurable neural computations rather than trial-and-error.

## Acknowledgements

We would like to acknowledge the Rice Wellbeing and Counseling Center, including Agnes Ho, as well as the administrative team in the Psychological Sciences Department at Rice University for help with participant recruitment. We thank Alexis Bailey and Ashwathi Nair for their assistance in participant recruitment and data collection.

## Author contributions

M.C.: Writing – original draft, Project administration, Methodology, Investigation, Formal analysis, Conceptualization. H.B.: Investigation, Data curation. L.F.: Investigation, Data curation. S.L.L: Writing – review & editing, Supervision, Methodology, Conceptualization. All authors reviewed and revised the final manuscript.

## Supplemental Methods

**Table S1.** Medicated vs. Unmedicated Participant Demographics.

| Variable | Responders<br>Mean (*SD*) | UMC - R<br>Mean (*SD*) | Non-Responders<br>Mean (*SD*) | UMC - NR<br>Mean (*SD*) |
| --- | --- | --- | --- | --- |
| *N* | 40 | 40 | 17 | 17 |
| Age* | 23.4 (4.96) | 20.5 (3.14) | 22.2 (2.92) | 20.4 (3.80) |
| Sex | 27F, 6M, 5N,<br>2T | 24F, 14M, 2N | 14F, 2M, 1N | 12F, 4M, 1N |
| Race |  |  |  |  |
| - White | 65% | 40% | 59% | 35% |
| - Asian/Asian<br>American | 30% | 55% | 29% | 53% |
| - Black/African<br>American | 10% | 10% | 24% | 0% |
| - American<br>Indian/Alaska<br>Native/Indigenous | 0% | 0% | 12% | 0% |
| - Arab/Middle<br>Eastern/North<br>African | 0% | 8% | 0% | 6% |
| - Native<br>Hawaiian/Pacific<br>Islander | 0% | 3% | 0% | 0% |
| - Multiracial | 8% | 3% | 12% | 12% |
| - Other | 3% | 3% | 0% | 12% |
| Ethnicity |  |  |  |  |
| - Hispanic/Latinx/a/o | 20% | 12% | 23% | 23% |
| Education (years<br>completed)*<sup>+</sup> | 15.5 (2.7) | 14.1 (2.0) | 15.5 (2.3) | 13.9 (1.7) |
| **Questionnaires** |  |  |  |  |
| BDI-II (current) | 9.4 (6.8) | 10.6 (7.1) | 22.8 (8.5) | 22.7 (8.8) |
| BAI (current)* | 12.2 (9.5) | 8.1 (7.1) | 17.1 (12.2) | 11.9 (6.7) |
| PHQ-9 | 5.4 (3.5) | 6.1 (4.7) | 9.9 (5.8) | 10.9 (5.7) |
| MDQ | 5.0 (4.0) | 4.3 (3.8) | 5.6 (3.2) | 6.1 (4.0) |
| ASRMS | 2.9 (3.2) | 3.0 (3.0) | 2.5 (2.7) | 3.0 (2.5) |
| **Neuropsychiatric Measures** |  |  |  |  |
| Perceived Stress Scale (past<br>month) | 15.8 (7.0) | 16.7 (5.5) | 21.4 (4.8) | 20.8 (4.9) |
| Perceived stress scale<br>(current)*<sup>+</sup> | 2.8 (2.6) | 3.7 (2.8) | 6.0 (2.5) | 5.4 (2.0) |
| Hours of sleep (last night) | 7.6 (3.4) | 7.1 (1.4) | 6.4 (1.9) | 7.1 (1.3) |
| Hours or sleep (past month) | 7.3 (1.1) | 6.9 (1.3) | 6.8 (1.3) | 7.0 (1.6) |
| ASRS | 10.7 (4.0) | 9.8 (3.7) | 12.0 (3.9) | 12.6 (3.0) |
| Y-BOCS | 7.8 (6.9) | 7.8 (6.3) | 7.8 (4.8) | 10.7 (7.3) |
| **Behavioral Task Accuracy** |  |  |  |  |
| Negative lure CR | 0.68 (0.11) | 0.74 (0.10) | 0.70 (0.14) | 0.73 (0.08) |
| Neutral lure CR | 0.77 (0.12) | 0.81 (0.09) | 0.78 (0.11) | 0.80 (0.10) |
| Positive lure CR | 0.66 (0.14) | 0.72 (0.13) | 0.70 (0.13) | 0.67 (0.12) |
| Negative lure FA | 0.32 (0.11) | 0.26 (0.10) | 0.30 (0.14) | 0.27 (0.08) |
| Neutral lure FA | 0.23 (0.12) | 0.19 (0.09) | 0.22 (0.11) | 0.20 (0.10) |
| Positive lure FA | 0.34 (0.14) | 0.28 (0.13) | 0.30 (0.13) | 0.33 (0.12) |

**Key:** ***BDI-II:** Beck Depressive Inventory II; **BAI:** Beck Anxiety Inventory; **PHQ-9:** Patient Health Questionnaire; **MDQ:** Mood Disorder Questionnaire; **ASRMS:** Altman Self-Rating Mania Scale; **ASRS:** The Adult ADHD Self-Report Scale (> 14 suggests presence of ADHD); **Y-BOCS:** Yale-Brown Obsessive Compulsive Scale (0-40, larger total scores indicate greater symptom severity). M = Male, F = Female, NB = Non-binary, T = Transgender. Other = “none of the above”. \*Significant differences between R and UMC-R groups; <sup>+</sup>Significant differences between NR and UMC-NR groups.*

### Questionnaires

We administered a battery of questionnaires to assess depression severity, comorbid psychiatric symptoms, medical and treatment history, and other relevant factors. Participants completed a demographics questionnaire to characterize age, sex, race, ethnicity, and years of education. Additionally, we administered a neuropsychiatric history and an antidepressant usage questionnaire (see Supplement for full questionnaire) to ascertain past depressive symptoms, behavioral therapy, and the duration and type of antidepressant use. Other questionnaires to assess neuropsychiatric symptoms included the PHQ-9 (1) to assess depressive symptoms and the BAI (2) to measure symptoms of anxiety. We also collected retrospective anxiety symptoms using the BAI with similar modified instructions to the BDI-II. We included this because anxiety symptoms are often highly comorbid with depressive symptoms; however, we were primarily interested in antidepressants’ influence on depressive symptoms for the purposes of the current study. We used the Mood Disorders Questionnaire (MDQ) (3) for bipolar disorder symptoms and the Altman Self-Rating Mania Scale (4) for mania symptoms. Perceived stress was measured with the Perceived Stress Scale (PSS) (5), and sleep health was evaluated using the Pittsburgh Sleep Quality Index (PSQI) (6). Other self-report measures included the Adult Attention-Deficit/Hyperactivity Disorder (ADHD) Self-Report Screening Scale (ASRS-5) (7) for attention deficits and the Yale Brown Obsessive Compulsive Scale (Y-BOCS) (8) for Obsessive-Compulsive Disorder (OCD) symptoms. Additionally, participants completed a Reproductive Survey covering menstrual health, birth control, and pregnancy history. These additional questionnaires were not the focus of the current study but were not included when determining if there were group differences across responder status and mechanism.

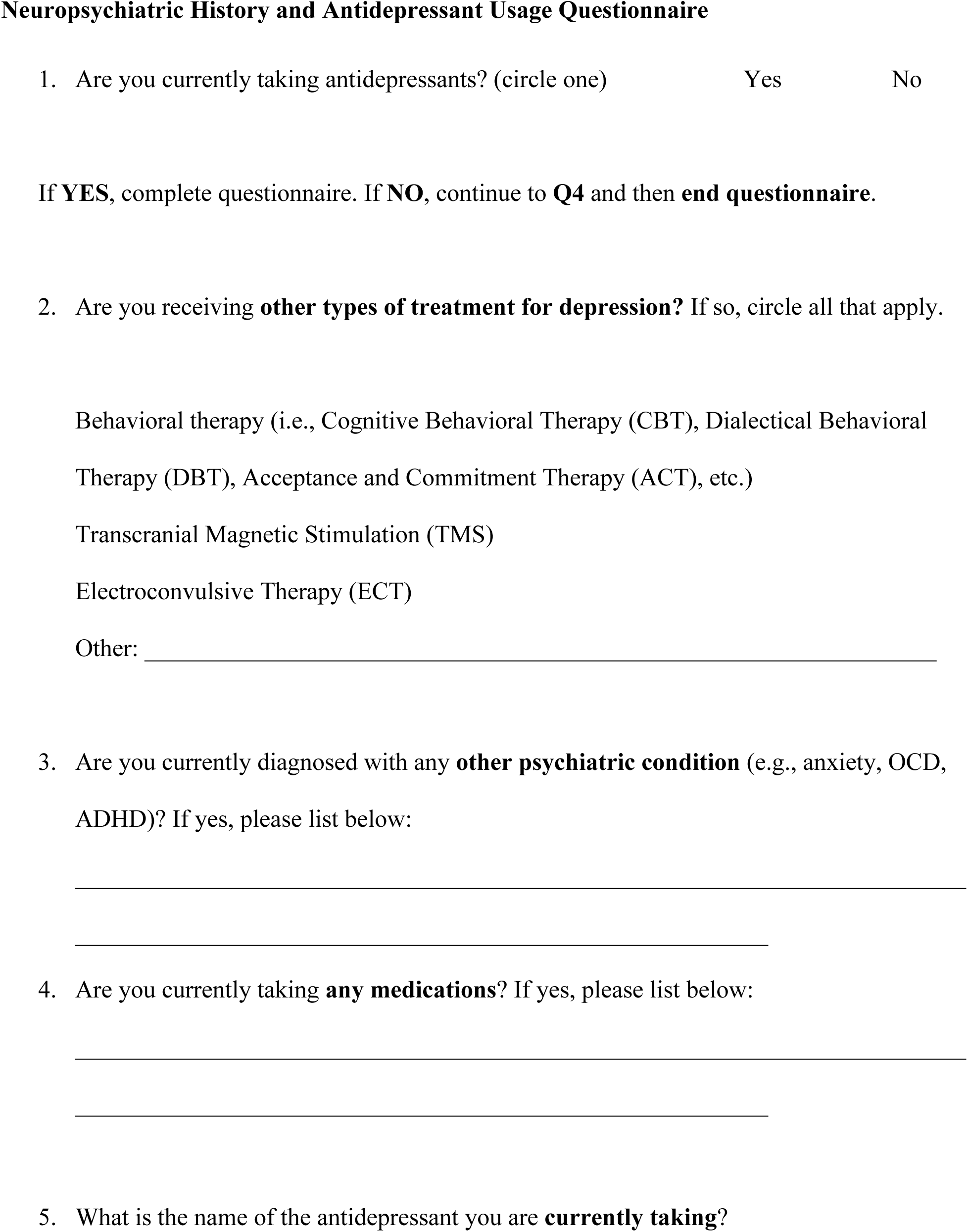
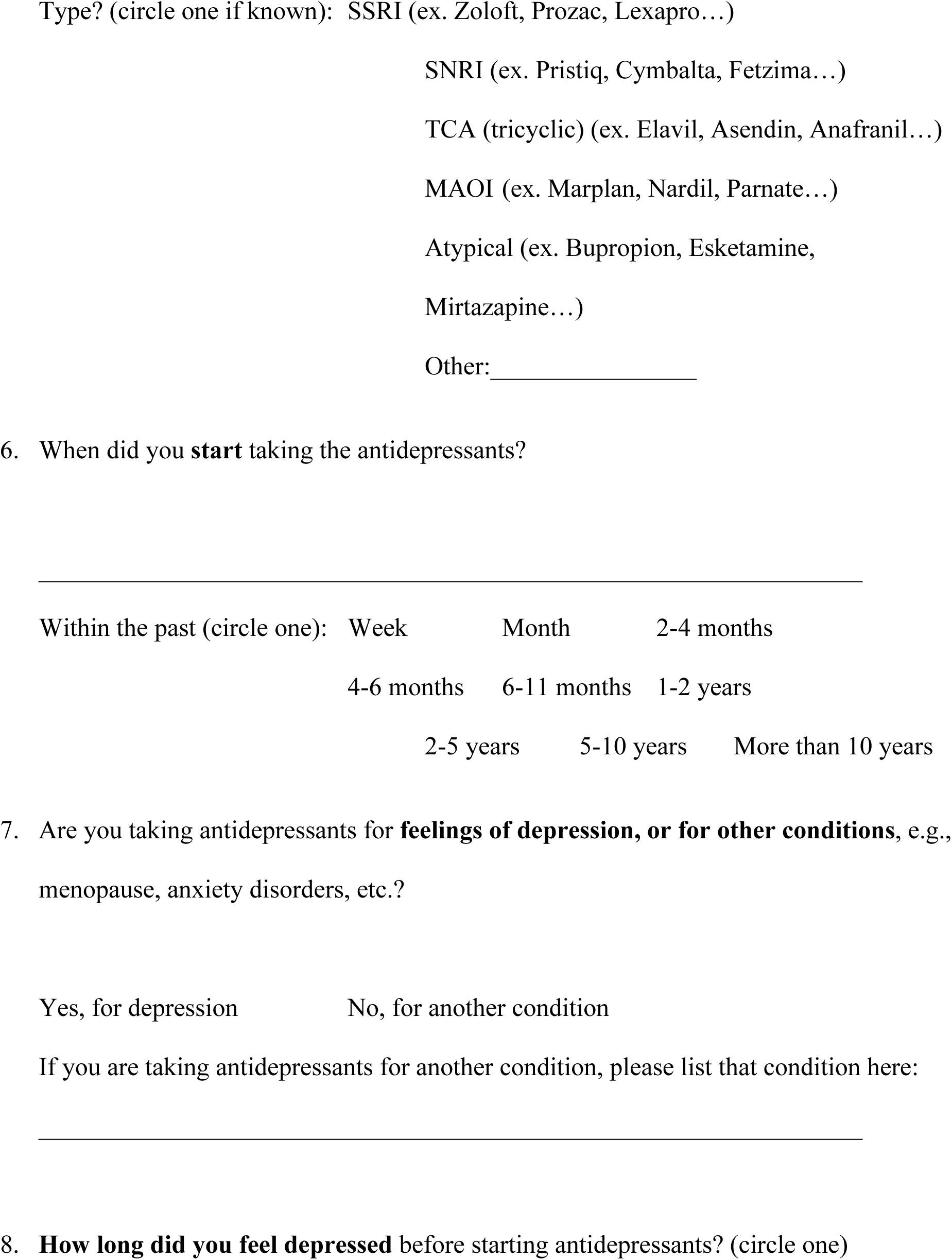
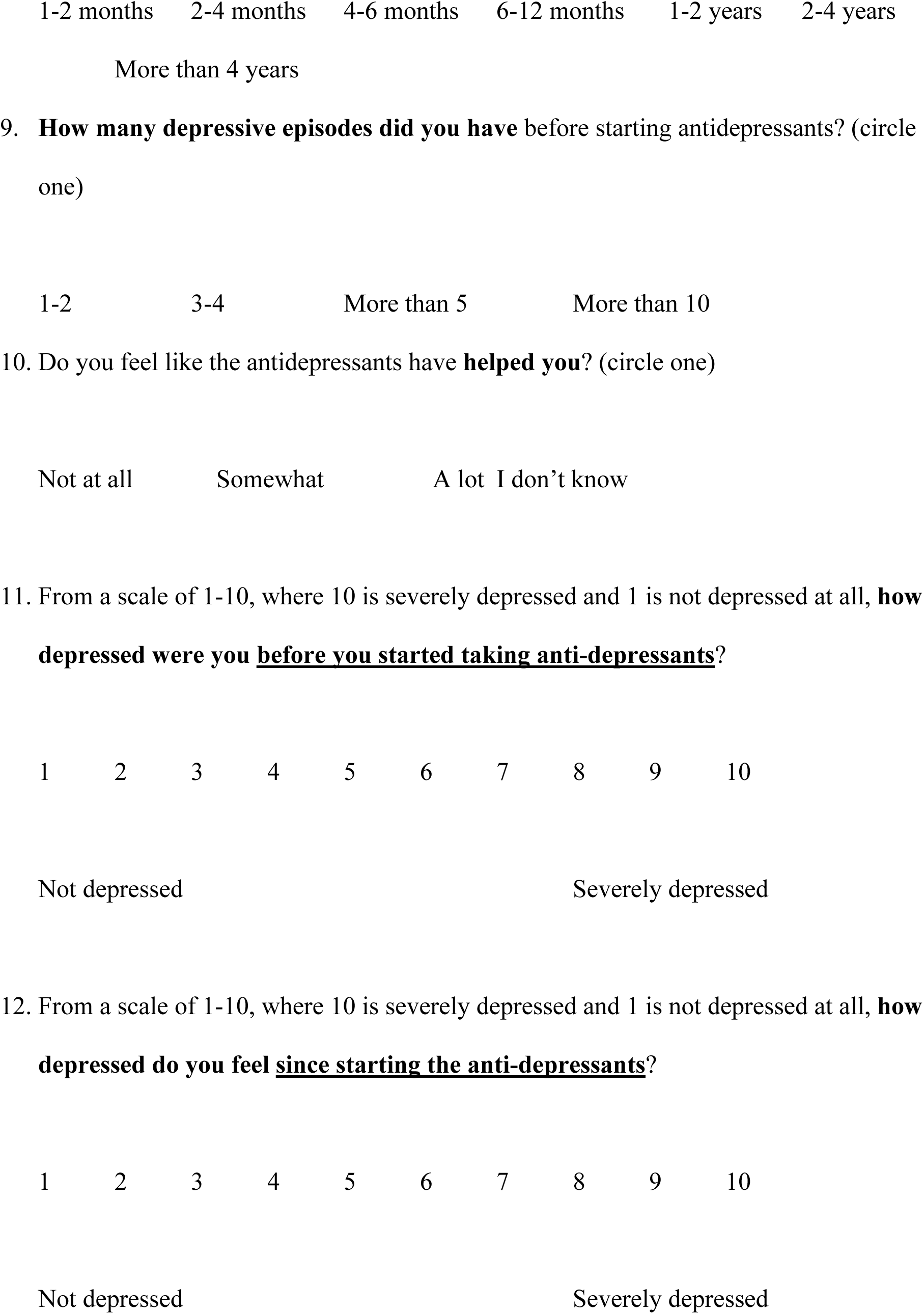
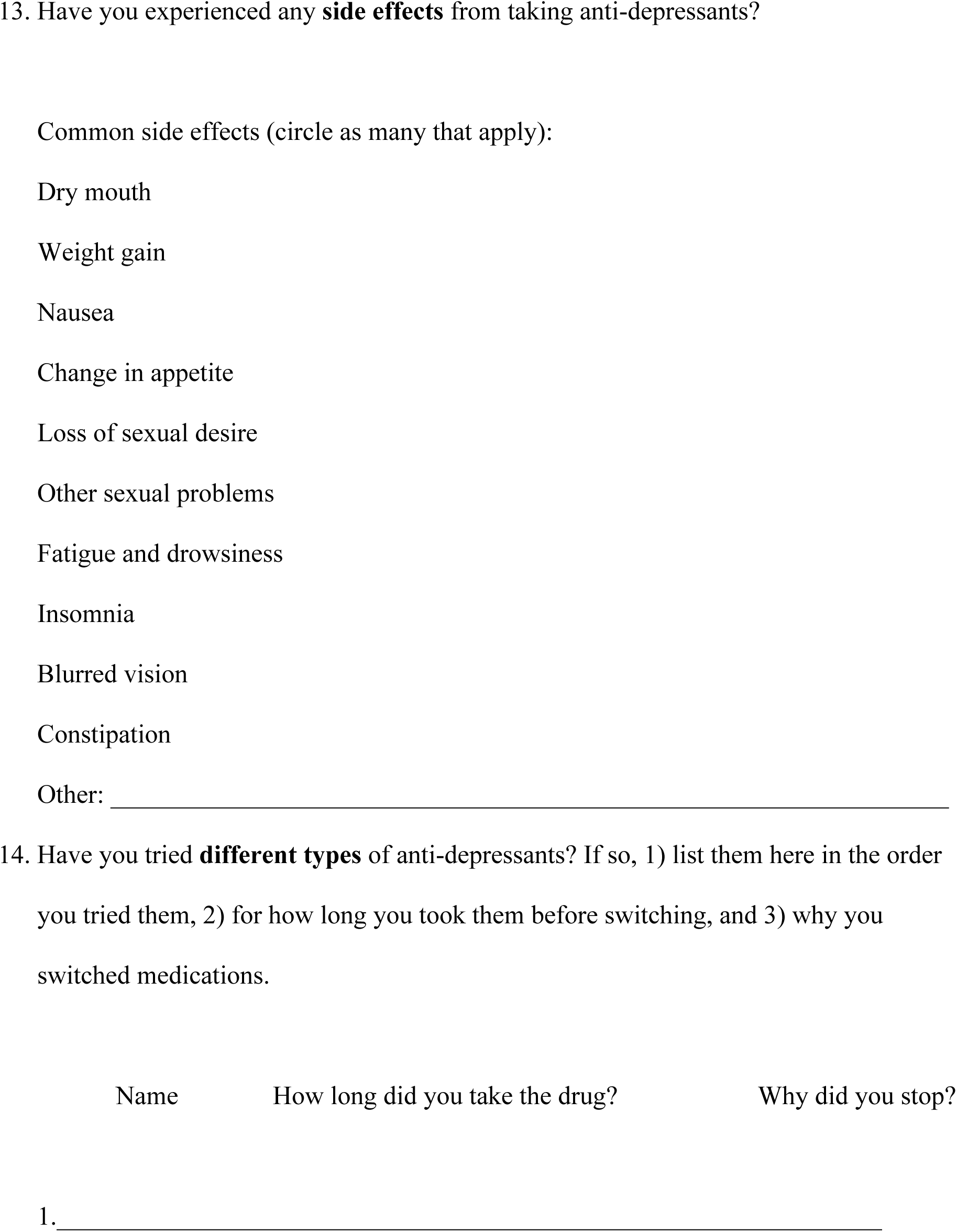
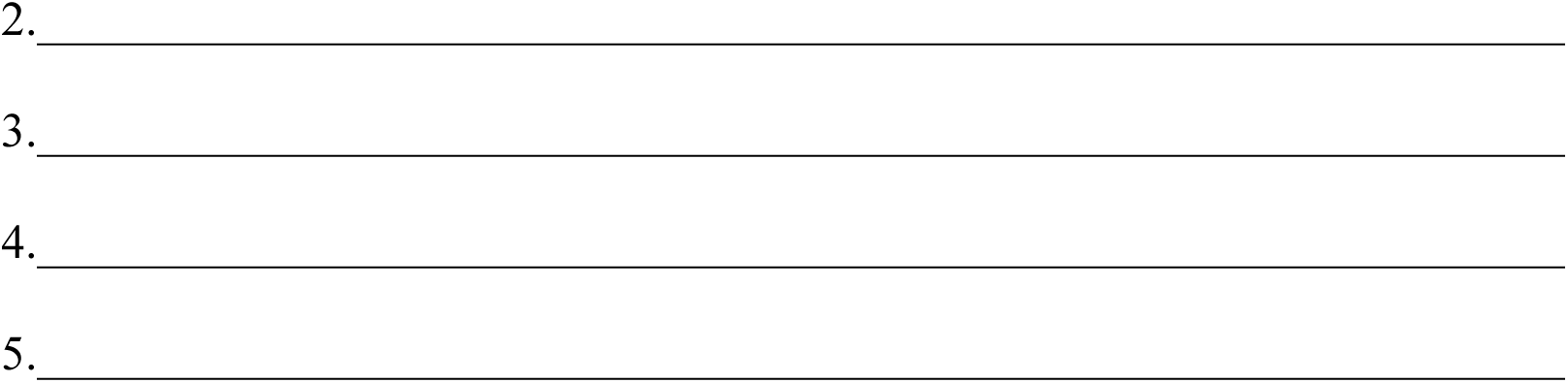

## Supplemental Results

### Supplemental Results 1

#### Group differences in demographic, neuropsychiatric, and psychological symptom measures across responders, non-responders, and matched unmedicated controls

First, independent samples t-tests were conducted to determine potential differences in demographic and questionnaire data between R and UMC - R. Variables included age, years of education, number of depressive episodes, number of times switching antidepressants, duration of antidepressant usage, BDI-II (current), BDI-II (retrospective), BAI (current), BAI (retrospective), symptoms of bipolar disorder (MDQ), symptoms of mania, average hours of sleep in the past month, average hours of sleep last night, current level of stress (PSS), average past month of stress (PSS), symptoms of OCD (Y-BOCS), and symptoms of ADHD (ASRS-5). We found differences in age [*t*(78) = 3.07, *p* = .003, *d* = 0.69], years of education [*t*(78) = 2.66, *p* = .010, *d* = 0.59], and BAI (current) [*t*(78) = 2.19, *p* = .031, *d* = 0.49], with R reporting greater age, education, and anxiety. Next, we conducted the same independent samples t-tests between NR and matched UMC - NR. We found differences in years of education [*t*(32) = 2.34, *p* = .025, *d* = 0.80], with NR reporting greater years of education.

### Supplemental Results 2

**Figure S2.**
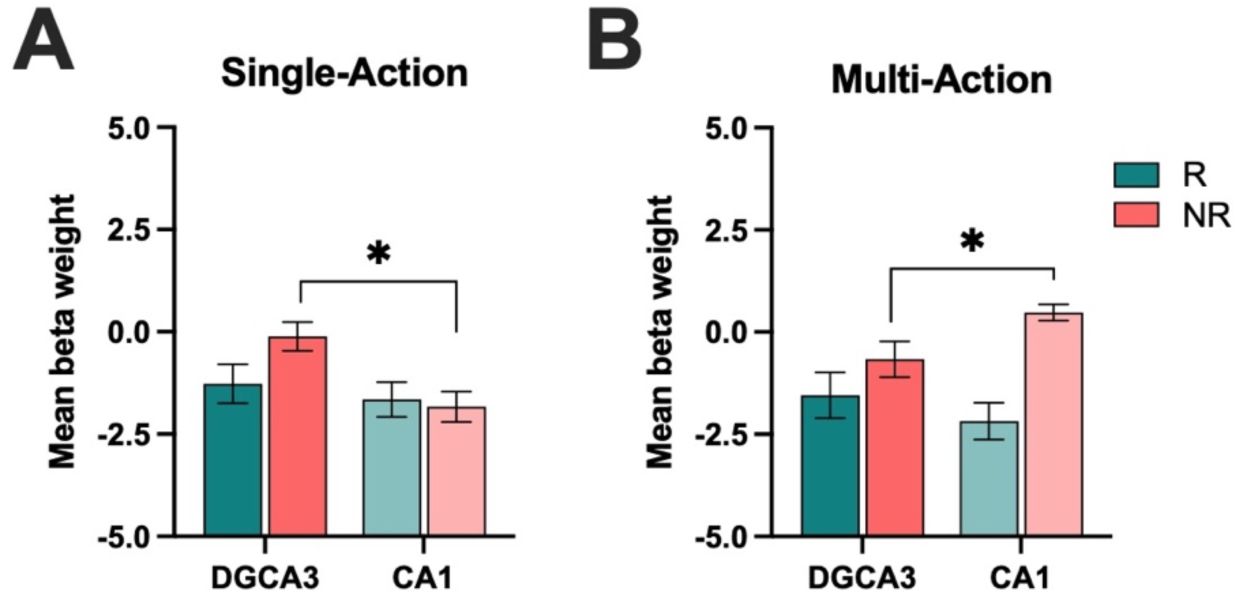
A) Mean activity beta weights during lure CRs in the DG/CA3 and CA1 across responder status in the single-action antidepressant group. B) Mean activity beta weights during lure CRs in the DG/CA3 and CA1 across responder status in the multi-action antidepressant group (*p < .05 and represents three-way interaction between region, mechanism, and responder status).

#### Independent models: responder-only and mechanism-only (hippocampus)

We conducted a mixed ANOVA on lure CR activity with emotion (Negative, Neutral, Positive) and region (DG/CA3, CA1) as within-subject factors and responder status (R, NR) as a between-subjects factor. There was a main effect of region [*F*(1, 67) = 5.47, *p* = .022, 17^2^ = 0.005; Fig. 2A], with greater DG/CA3 activity compared to CA1. There were no other significant main effects or interactions. Next, we tested the independent effect of mechanism (single-action,multi-action) on hippocampal (DG/CA3, CA1) activity during negative, neutral, and positive lure CRs. We conducted a mixed ANOVA on lure CR activity with emotion (Negative, Neutral, Positive) and region (DG/CA3, CA1) as within-subject factors and mechanism (single-action, multi-action) as a between-subjects factor. Similarly, we found a main effect of region [*F*(1, 67) = 4.52, *p* = .037, 17^2^ = 0.004], with greater DG/CA3 activity compared to CA1. There were no other main effects or interactions (Fig. 2B).

#### Posterior hippocampal models and full models with axis

We conducted a mixed ANOVA on lure CR activity with emotion (Negative, Neutral, Positive) and region (DG/CA3, CA1) as within-subject factors and both responder status (R, NR) and mechanism (single-action, multi-action) as between-subjects factors in the posterior hippocampus. There were no significant main effects or interactions [*p* > .05] (Fig. S3A-B. However, when we included hippocampal axis as a within subjects’ factor (anterior, posterior), we found a significant emotion x region x mechanism interaction [*F*(2, 124) = 3.52, *p* = .034, 17^2^ = 0.054], where DG/CA3 activity was greater than CA1 activity for emotional (negative and positive) relative to neutral lure CRs and was selective to those taking SSRIs relative to multi-action antidepressants [F(1,62) = 7.70, p = .007, 17^2^ = 0.110] (Fig S3C-D). There were also marginal effects of axis [F(1,62) = 3.18, p = .08, 17^2^ = 0.049] and axis x emotion [F(2,124) = 2.80, p = .065, 17^2^ = 0.043], supporting that effects were selective to anterior relative to posterior hippocampus.

**Figure S3.**
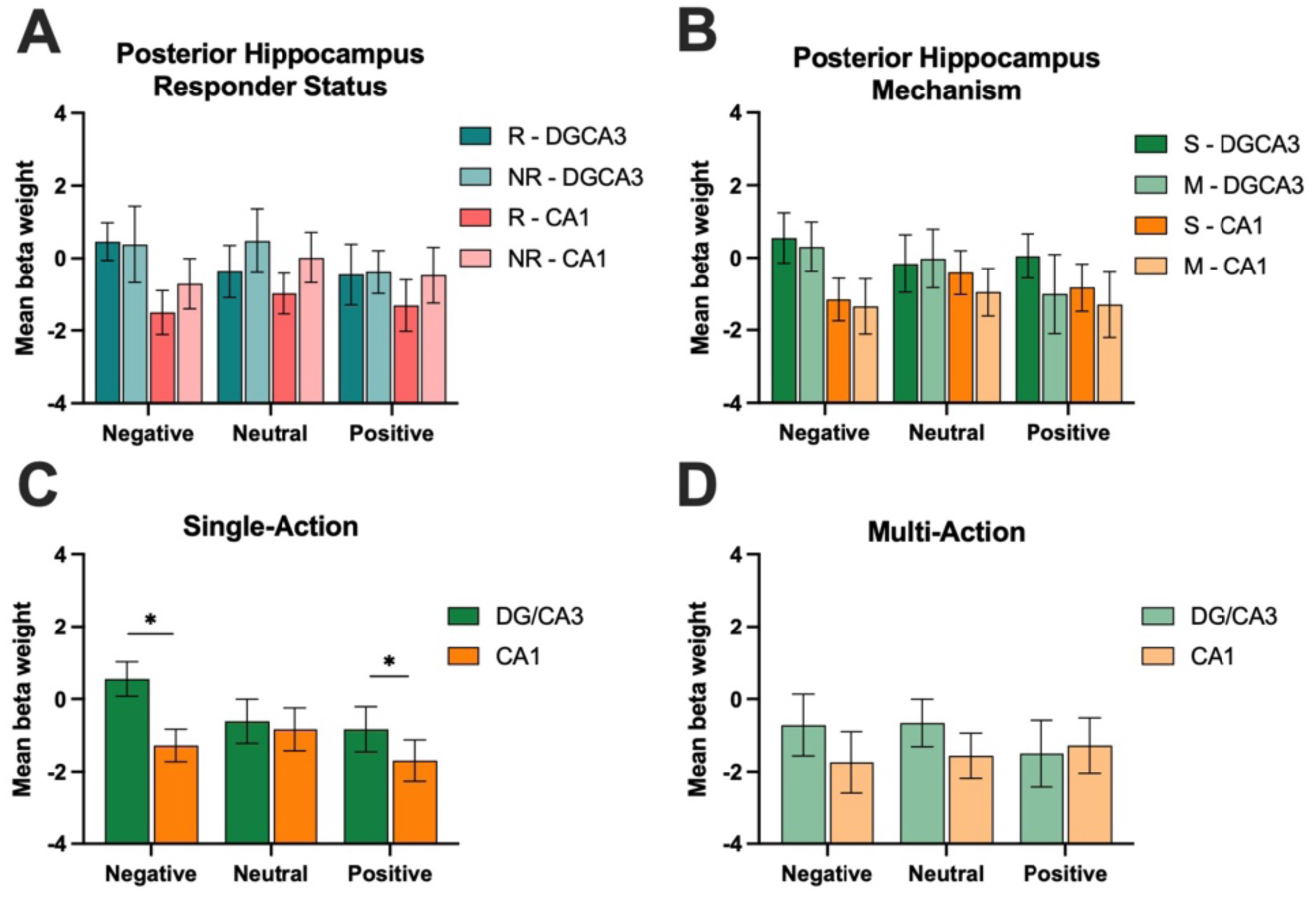
Posterior hippocampal DG/CA3 and CA1 lure CR activity across responder status and mechanism. A) Mean activity beta weights during lure CRs across responder status; B) Mean activity beta weights during lure CRs across mechanism. C) Mean whole hippocampal activity beta weights during lure CRs across single-action and D) multi-action groups (*p < .05 and represents an emotion by region by mechanism interaction).

### Supplemental Results 3

#### Responder status and mechanism similarly impact hippocampal subfield activity during lure FAs

We conducted a mixed ANOVA on lure FA activity with emotion (Negative, Neutral, Positive) and region (DG/CA3, CA1) as within-subject factors, and both responder status (R, NR) and mechanism (single-action, multi-action) as between-subjects factors. We found a main effect of region [*F*(1, 64) = 6.62, *p* = .012, 17^2^ = 0.008], where there was greater activity in the DG/CA3 compared to the CA1. Next, we performed the exploratory models independently by responder status and mechanism. We conducted a mixed ANOVA on lure FA activity with emotion (Negative, Neutral, Positive) and region (DG/CA3, CA1) as within-subject factors, and responder status (R, NR) as a between-subjects factor. There was a main effect of emotion [*F*(2, 113) = 3.74, *p* = .033, 17^2^ = 0.023], such that there was greater activity during positive lure FAs compared to negative or neutral [*F*(1,66) = 5.42, *p* = .023, 17^2^ = 0.076]. There was also a main effect of region [*F*(1, 66) = 6.32, *p* = .014, 17^2^ = 0.007], where there was greater activity during lure FAs in the DG/CA3 compared to the CA1. There were no other main effects or interactions (Fig. S4A).

**Figure S4.**
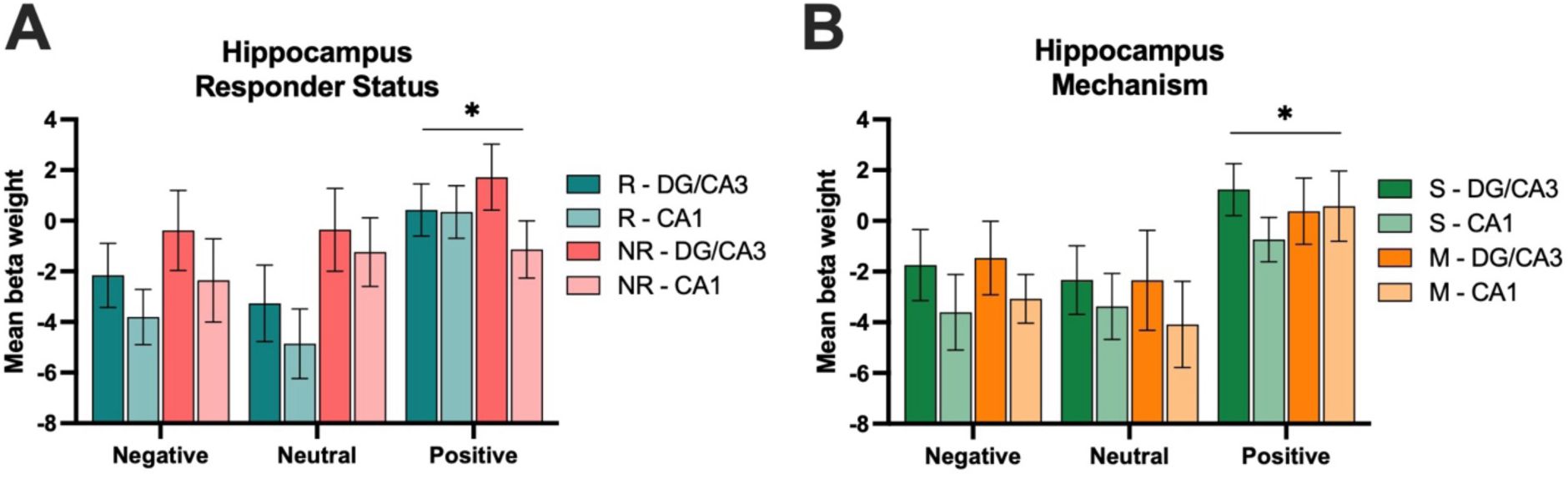
Hippocampal DG/CA3 and CA1 lure FA activity across responder status and mechanism. A) Mean activity beta weights during lure FAs across responder status; B) Mean activity beta weights during lure FAs across mechanism (*p < .05).

Next, we conducted the same analyses within mechanism groups. We conducted a mixed ANOVA on lure FA activity with emotion (Negative, Neutral, Positive) and region (DG/CA3, CA1) as within-subject factors, and mechanism (single-action, multi-action) as a between-subjects factor. Similar to responder status, we found a main effect of emotion [*F*(2, 114) = 5.84, *p* = .006, 17^2^ = 0.036], where there was greater activity during positive lure FAs compared to negative or neutral [*F*(1,66) = 7.58, *p* = .008, 17^2^= 0.103]. There was also a main effect of effect of region [*F*(1, 66) = 5.87, *p* = .018, 17^2^ = 0.007], where there was greater activity during lure FAs in the DG/CA3 compared to the CA1. We found no other main effects or interactions (Fig. S4B).

#### Minimal effects of responder status, but not mechanism, on lure FAs in posterior hippocampal regions

We followed the same framework for lure FAs across responder status and mechanism in the posterior hippocampus. We conducted a mixed ANOVA on lure FA activity with emotion (Negative, Neutral, Positive) and posterior region (DG/CA3, CA1) as within-subject factors, and both responder status (R, NR) and mechanism (single-action, multi-action) as between-subjects factors. We found a marginally significant emotion x region x responder status interaction [*F*(2, 122) = 2.94, *p* = .057, 17^2^ = 0.009] (Fig. S5). To formally assess axis specificity, we extended the model to include hippocampal axis (anterior vs. posterior) as a within-subjects factor. In the full model including both responder status and mechanism, there was a main effect of region [*F*(1, 61) = 7.44, *p* = .008, 17^2^ = 0.005]. There were no other significant main effects or interaction [*p* > .05].

**Figure S5.**
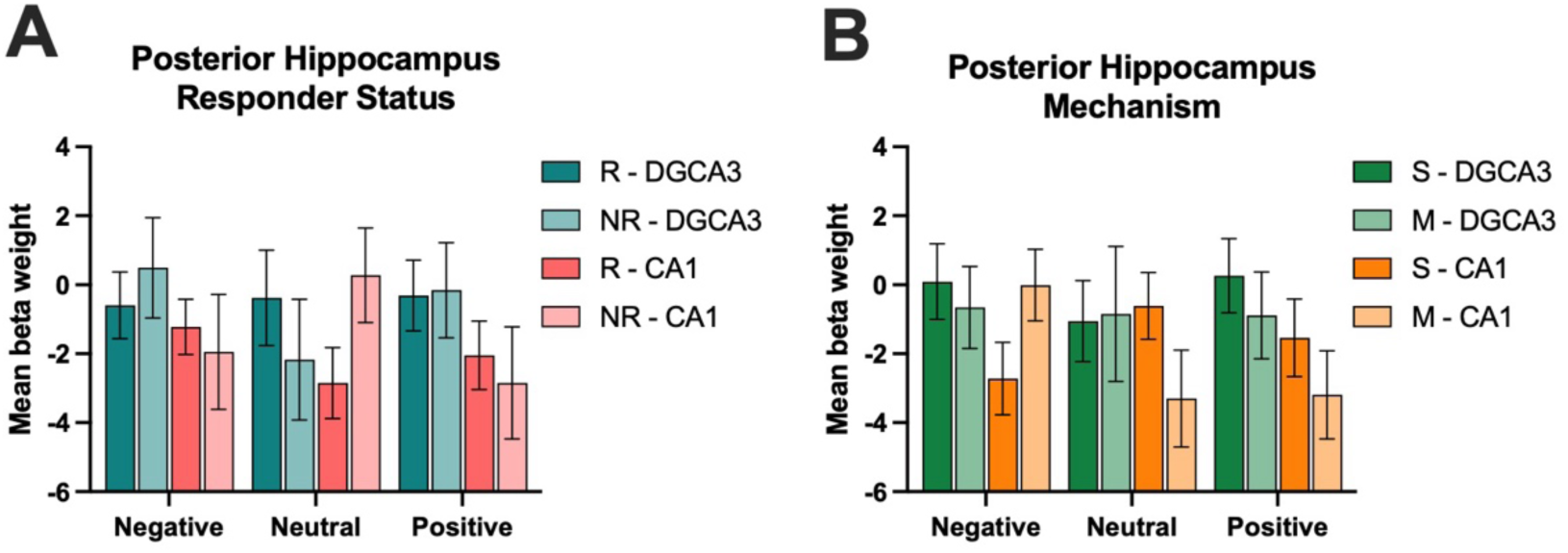
Posterior hippocampal DG/CA3 and CA1 lure FA activity across responder status and mechanism. A) Mean activity beta weights during lure FAs across responder status; B) Mean activity beta weights during lure FAs across mechanism (*p < .05).

### Supplemental Results 4

**Figure S6.**
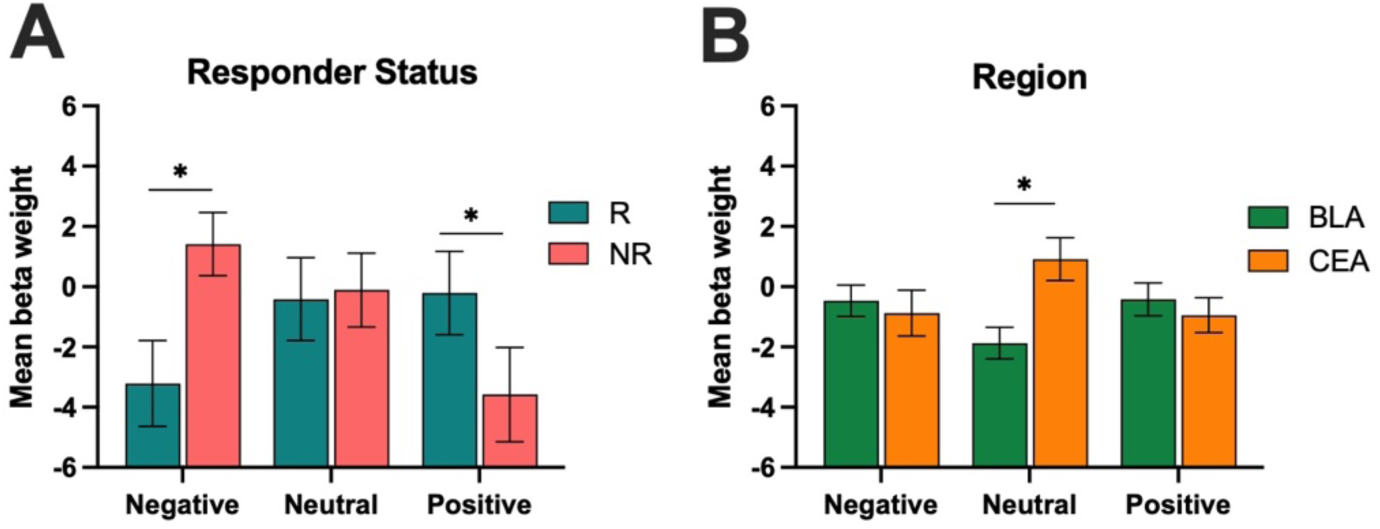
Visualization of interactions in amygdala models. A) Mean activity beta weights during lure CRs in R and NR collapsed across amygdala subnuclei. B) Mean activity beta weights during lure CRs in the BLA and CEA collapsed across responder status (*p < .05 and represents emotion x responder status and emotion x region interaction).

#### Independent models: responder-only and mechanism-only (amygdala)

We conducted a mixed ANOVA on lure CR activity with emotion (Negative, Neutral, Positive) and region (BLA, CEA) as within-subject factors and responder status (R, NR) as a between-subjects factor. There was a significant interaction between emotion and responder status [*F*(2, 134) = 4.47, *p* = .013, 17^2^ = 0.021], where R had lower amygdala activity during negative lure CRs compared to NR but greater amygdala activity during positive lure CRs [*F*(1,67) = 7.26, *p* = .009, 17^2^ = 0.098]. There was also an interaction between emotion and region [*F*(2, 134) = 13.85, *p* < .001, 17^2^ = 0.022], where there was greater activity during neutral lure CRs in the CEA compared to the BLA, with no differences across region for emotional lure CRs [*F*(1,67) = 31.93, *p* < .001, 17^2^ = 0.323] (Fig. 2C). There were no other significant main effects or interactions.

Next, we tested the independent effect of mechanism (single-action, multi-action) on amygdala (BLA, CEA) activity during negative, neutral, and positive lure CRs. We conducted a mixed ANOVA on lure CR activity with emotion (Negative, Neutral, Positive) and region (BLA, CEA) as within-subject factors and mechanism (single-action, multi-action) as a between-subjects factor. There was a significant emotion x mechanism interaction [*F*(2, 134) = 3.13, *p* = .047, 17^2^ = 0.015], where those taking single-action antidepressants showed greater amygdala activity during negative lure CRs but lower amygdala activity during positive lure CRs compared to those taking multi-action antidepressants [*F*(1,67) = 40.53, *p* < .001, 17^2^ = .377]. There was also a significant emotion x region interaction [*F*(2, 134) = 18.00, *p* < .001, 17^2^ = 0.028], where there was greater CEA activity compared to BLA activity for neutral lure CRs, with no differences across region for emotional lure CRs [*F*(1,67) = 40.53, *p* < .001, 17^2^ = 0.377 (Fig. 2D). We found no other main effects or interactions in this model.

#### Supplemental Results 5

**Figure S7.**
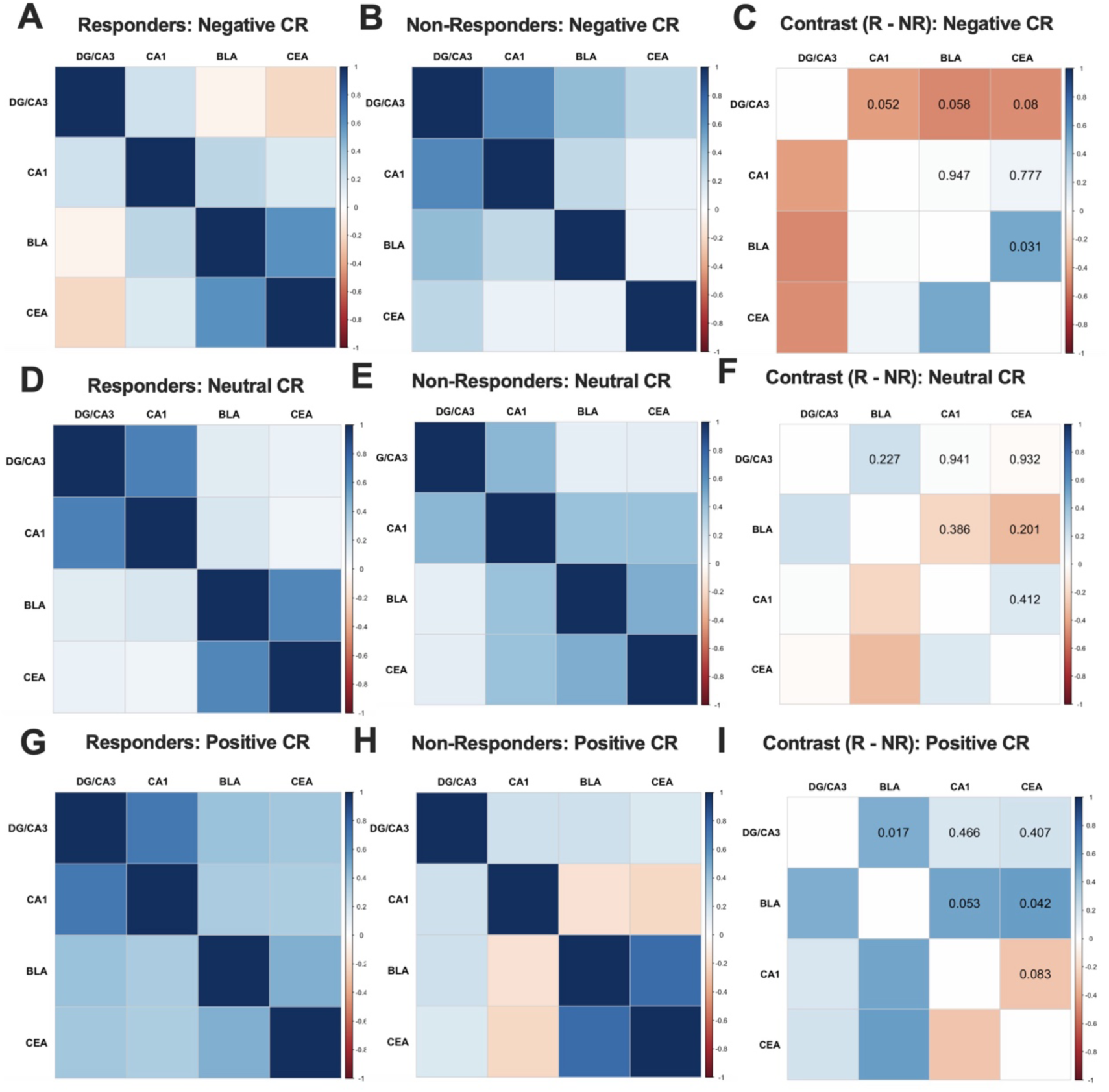
Posterior coactivation for lure CRs in R and NRs. A) Negative lure CRs correlation matrices for pairwise r^2^ values across posterior hippocampal (DG/CA3, CA1) and amygdala (BLA, CEA) ROIs in R and B) NR groups; C) Negative lure CRs contrast correlation matrix representing the difference in pairwise r values in R and NR; D) Neutral lure CRs correlation matrices for pairwise r values across posterior ROIs in R and E) NR; F) Neutral lure CRs contrast correlation matrix representing the difference in pairwise r values in R and NR; G) Positive lure CRs correlation matrices for pairwise r values across posterior ROIs in R and H) NR; I) Positive lure CRs contrast correlation matrix representing the difference in pairwise r values in R and NR. The contrast correlation matrices were normalized using Fisher’s r-to-z transformation and compared using a z-difference test (*p < .0083; Bonferroni corrected for multiple comparisons). Abbreviations: ROI, region of interest.

**Figure S8.**
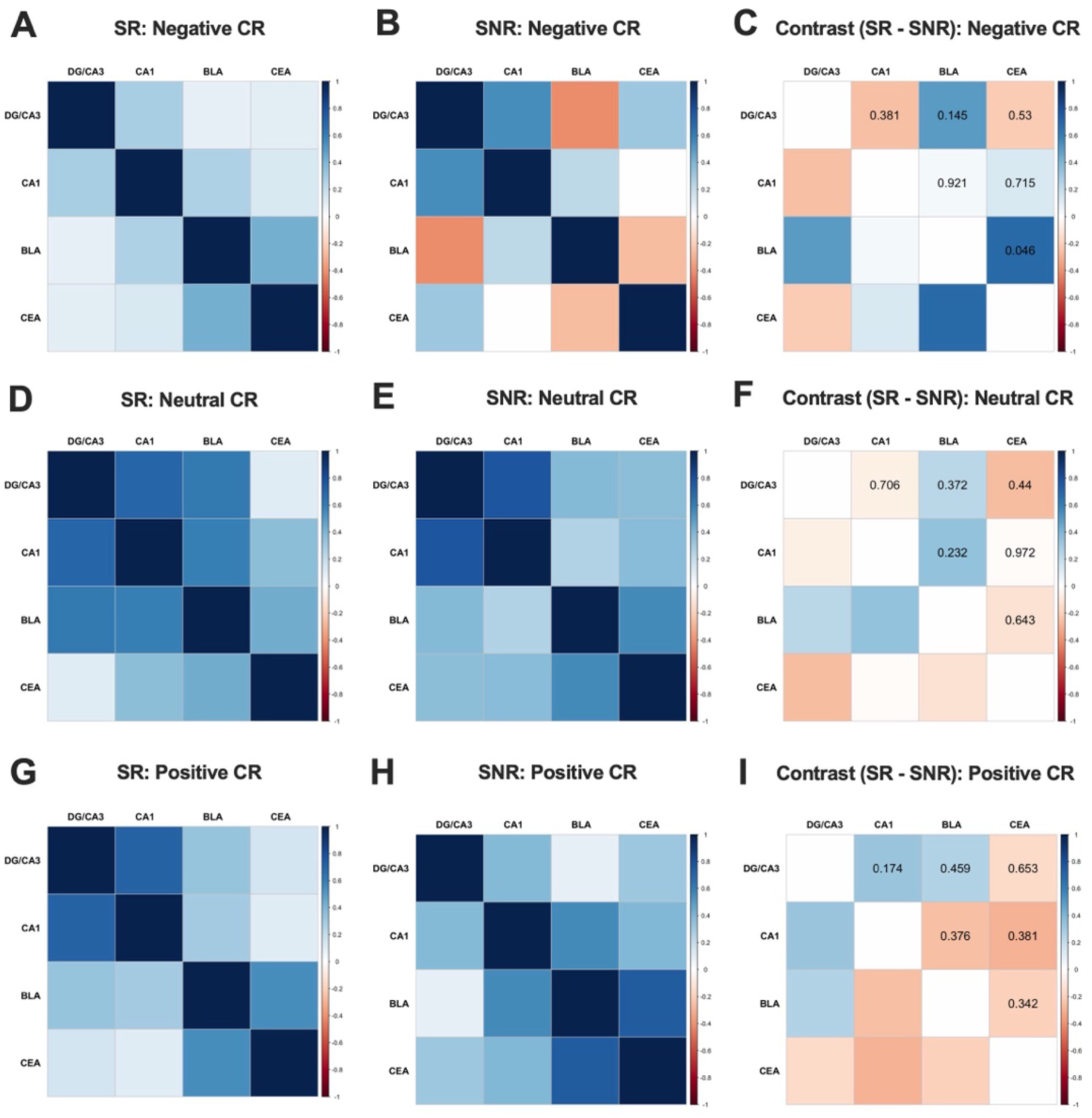
Functional coactivation for lure CRs in single-action R and NRs. A) Negative lure CRs correlation matrices for pairwise r values across ROIs in single-action R and B) NR. C) Negative lure CRs contrast correlation matrix representing the difference in pairwise r values in single-action R and NR. D) Neutral lure CRs correlation matrices for pairwise r values across ROIs in single-action R and E) NR. F) Neutral lure CRs contrast correlation matrix representing the difference in pairwise r values in single-action R and NR. G) Positive lure CRs correlation matrices for pairwise r values across ROIs in single-action R and H) NR. I) Positive lure CRs contrast correlation matrix representing the difference in pairwise r values in single-action R and NR. The contrast correlation matrices were normalized using Fisher’s r-to-z transformation and compared using a z-difference test (*p < .0083; Bonferroni corrected for multiple comparisons). Abbreviations: ROI, region of interest.

**Figure S9.**
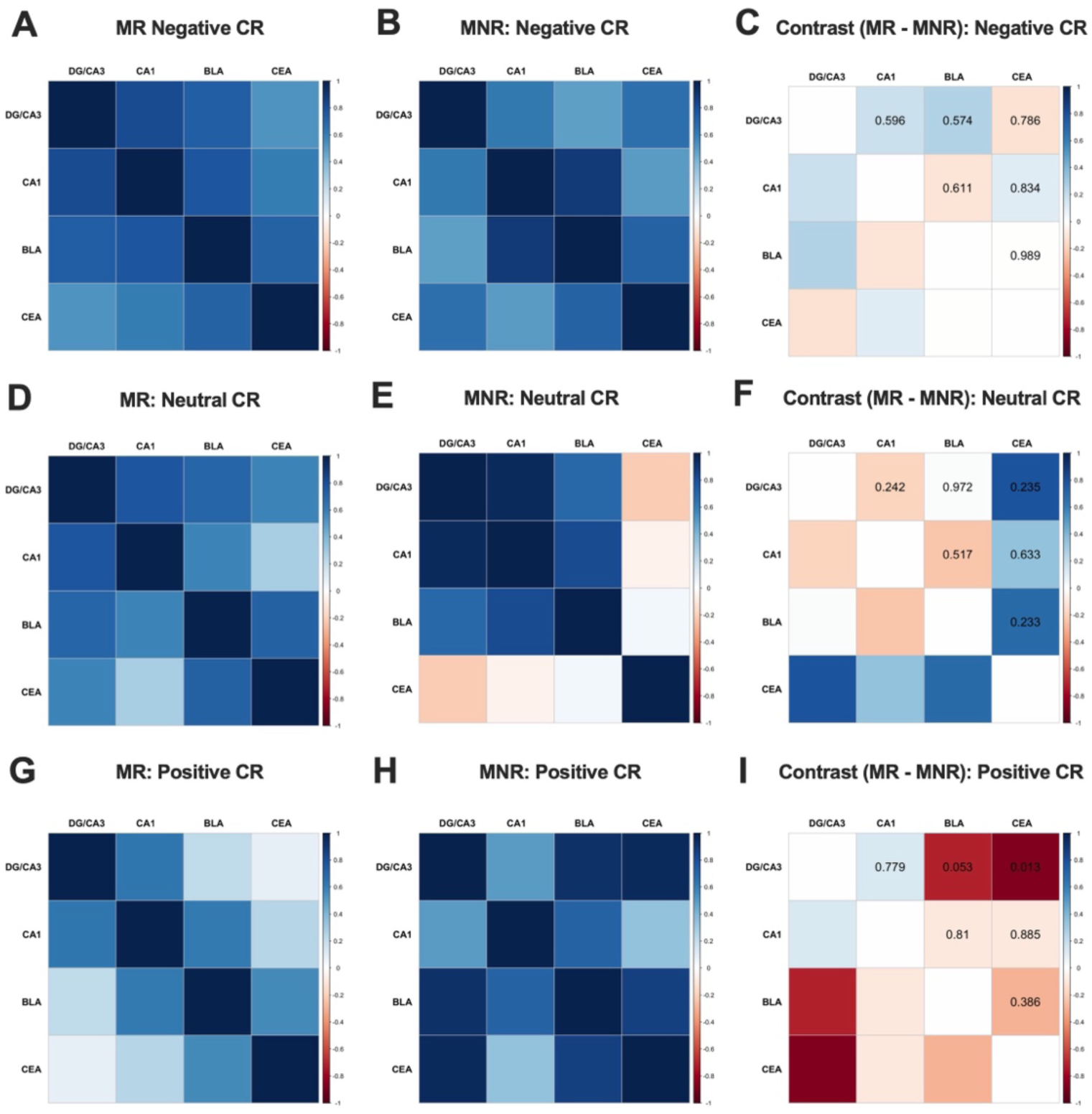
Functional coactivation for lure CRs in multi-action R and NRs. A) Negative lure CRs correlation matrices for pairwise r values across ROIs in multi-action R and B) NR. C) Negative lure CRs contrast correlation matrix representing the difference in pairwise r values i multi-action R and NR. D) Neutral lure CRs correlation matrices for pairwise r values across ROIs in multi-action R and E) NR. F) Neutral lure CRs contrast correlation matrix representing the difference in pairwise r values in multi-action R and NR. G) Positive lure CRs correlation matrices for pairwise r values across ROIs in multi-action R and H) NR. I) Positive lure CRs contrast correlation matrix representing the difference in pairwise r values in multi-action R and NR. The contrast correlation matrices were normalized using Fisher’s r-to-z transformation and compared using a z-difference test (*p < .0083; Bonferroni corrected for multiple comparisons). Abbreviations: ROI, region of interest.

Next, we examined whether hippocampal-amygdala functional coactivation differed as a function of mechanism group within the posterior hippocampal regions, which revealed no significant differences across mechanism (Fig. S10). However, when examining the whole hippocampus, we found that the multi-action mechanism group exhibited significantly greater DG/CA3-BLA (r = .630, p < .001) and DG/CA3-CA1 (*r* = .798, p < .001) functional coactivation during negative lure CRs compared to the single-action mechanism group (DG/CA3-BLA: *r* = 0.016, *p* = 0.925; DG/CA3-CA1: *r* = .330, *p* = 0.046) [DG/CA3-BLA: *z* = -2.87, *p* = .004; DG/CA3-CA1: *z* = −2.97, *p* = .003]

**Figure S10.**
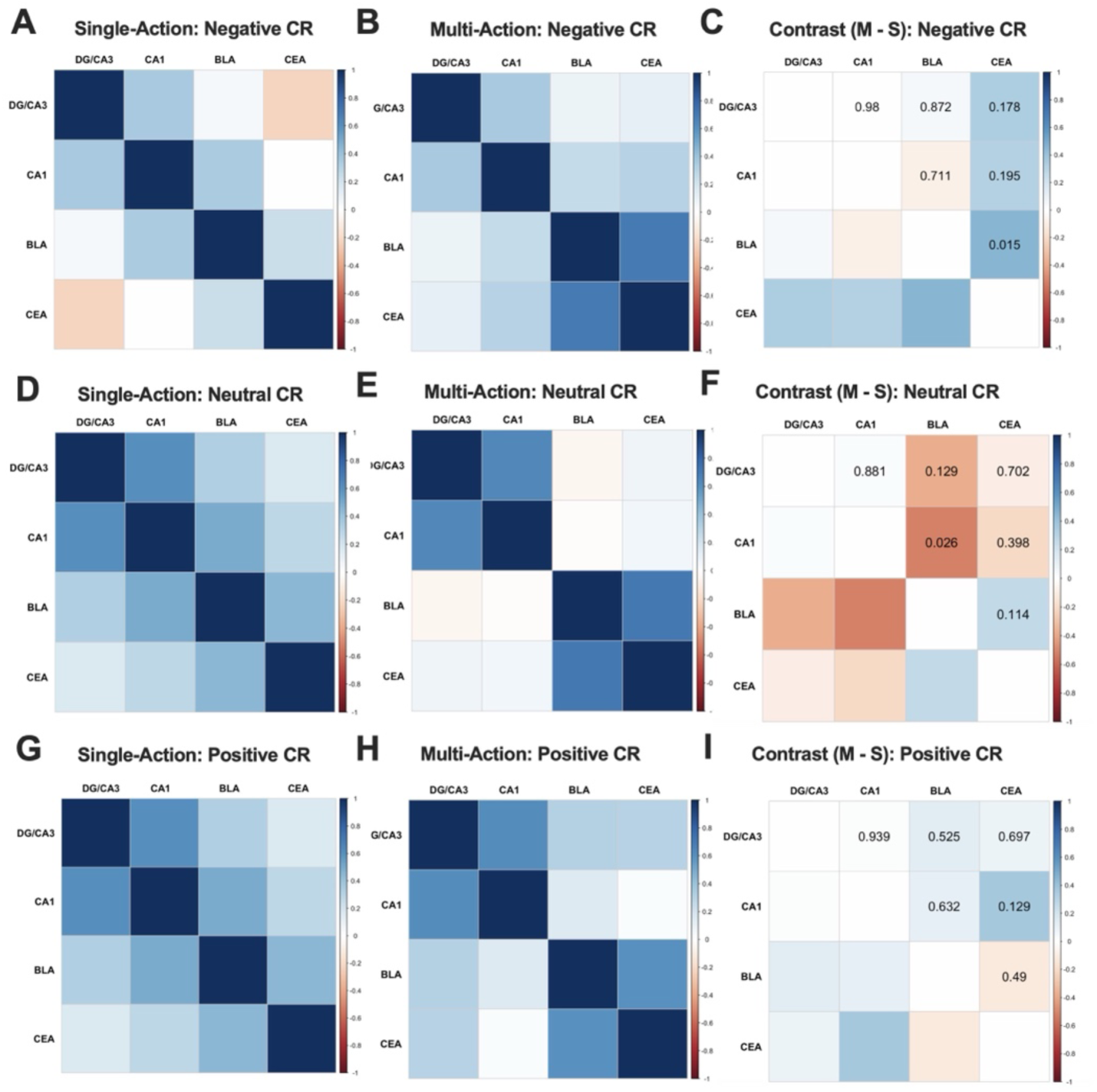
Posterior hippocampal-amygdala functional coactivation during lure CRs in single-action and multi-action mechanism groups. A) Negative lure CR correlation matrices for pairwise r values across posterior hippocampal (DG/CA3, CA1) and amygdala (BLA, CEA) regions of interest (ROIs) in single-action and B) multi-action groups; C) Negative lure CR contrast correlation matrix representing the difference in pairwise r values between single-action and multi-action groups. D) Neutral lure CR correlation matrices for pairwise r values across ROIs in single-action and E) multi-action groups; F) Neutral lure CR contrast correlation matrix representing the difference in pairwise r values between single-action and multi-action groups. G) Positive lure CRs correlation matrices for pairwise r values across ROIs in single-action and H) multi-action groups; I) Positive lure CRs contrast correlation matrix representing the difference in pairwise r values between single-action and multi-action groups. The contrast correlation matrices were normalized using Fisher’s r-to-z transformation and compared using a z-difference test (*p < .0083; Bonferroni corrected for multiple comparisons).

### Supplemental Results 6

#### Antidepressant mechanism, but not responder status, differentially impacts amygdala subnuclei during lure FAs

We ran the same set of analyses for lure FAs in the amygdala. We conducted a mixed ANOVA on lure FA activity with emotion (Negative, Neutral, Positive) and region (BLA, CEA) as within-subject factors, and both responder status (R, NR) and mechanism (Single, Multi) as between-subjects factors. We found no significant main effects or interactions. Next, we conducted a mixed ANOVA on lure FA activity with emotion (Negative, Neutral, Positive) and region (BLA, CEA) as within-subject factors, and responder status (R, NR) as a between-subjects factor. We found no significant main effects or interactions (Fig. S11A). Finally, we conducted the same analysis within mechanism groups. We conducted a mixed ANOVA on lure FA activity with emotion (Negative, Neutral, Positive) and region (BLA, CEA) as within-subject factors, and mechanism (single-action, multi-action) as a between-subjects factor. Here, we found an emotion x region interaction [*F*(2, 132) = 4.75, *p* = .010, 17^2^ = 0.011], where there was greater activity during positive lure FAs in the BLA compared to the CEA [*F*(1,66) = 6.70, *p* = .012, 17^2^ = 0.092]. There was also a three-way interaction between emotion, region, and mechanism [*F*(2, 132) = 3.38, *p* = .037, 17^2^ = 0.008], such that those taking multi-action antidepressants showed greater BLA activity during neutral lure FAs but lower CEA activity during positive lure FAs compared to those taking single-action antidepressants who showed little change in activity across emotion and region [*F*(1,66) = 5.80, *p* = .019, 17^2^ = 0.081]. We found no other main effects of interactions (Fig. S11B).

**Figure S11.**
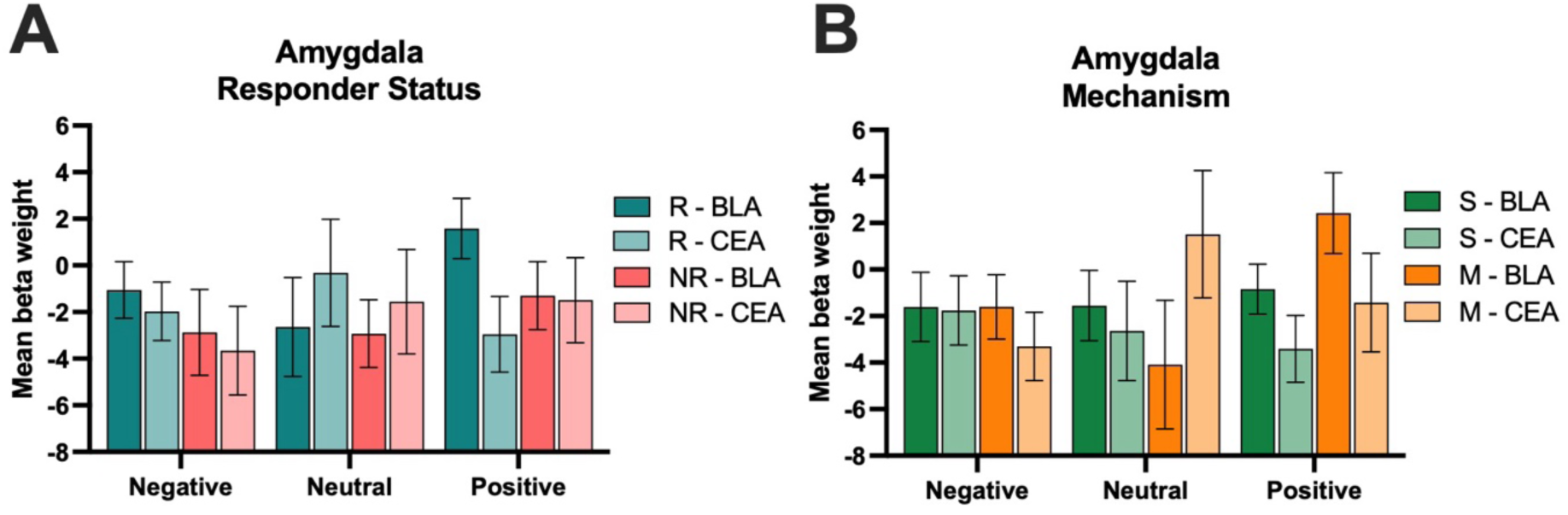
Amygdala BLA and CEA lure FA activity across responder status and mechanism. A) Mean activity beta weights during lure FAs across responder status; B) Mean activity beta weights during lure FAs across mechanism (*p < .05).

### Supplemental Results 7

#### No differences in hippocampal subfield activity between responders and unmedicated severity-matched controls

First, we conducted a mixed ANOVA on lure CR activity with emotion (Negative, Neutral, Positive) and region (DG/CA3, CA1) as within-subject factors and group (R, UMC-R) as a between-subjects factor in anterior hippocampus. There was a main effect of region [*F*(1, 78) = 11.45, *p* = .001, 17^2^ = 0.006], where there was greater DG/CA3 activity during lure CRs relative to CA1 (Fig. 5A). In an exploratory analysis, we also stratified the R by single-action and multi-action mechanism and compared to the UMC-R group. We conducted a mixed ANOVA on lure CR activity with emotion (Negative, Neutral, Positive) and region (DG/CA3, CA1) as within-subject factors, and group (single-action R, multi-action R, UMC-R) as a between-subjects factor. There was a main effect of region [*F*(1, 77) = 7.28, *p* = .009, 17^2^ = 0.00], where there was greater DG/CA3 activity relative to CA1 (Fig. S12A-B), but no main effects or interactions with group.

**Figure S12.**
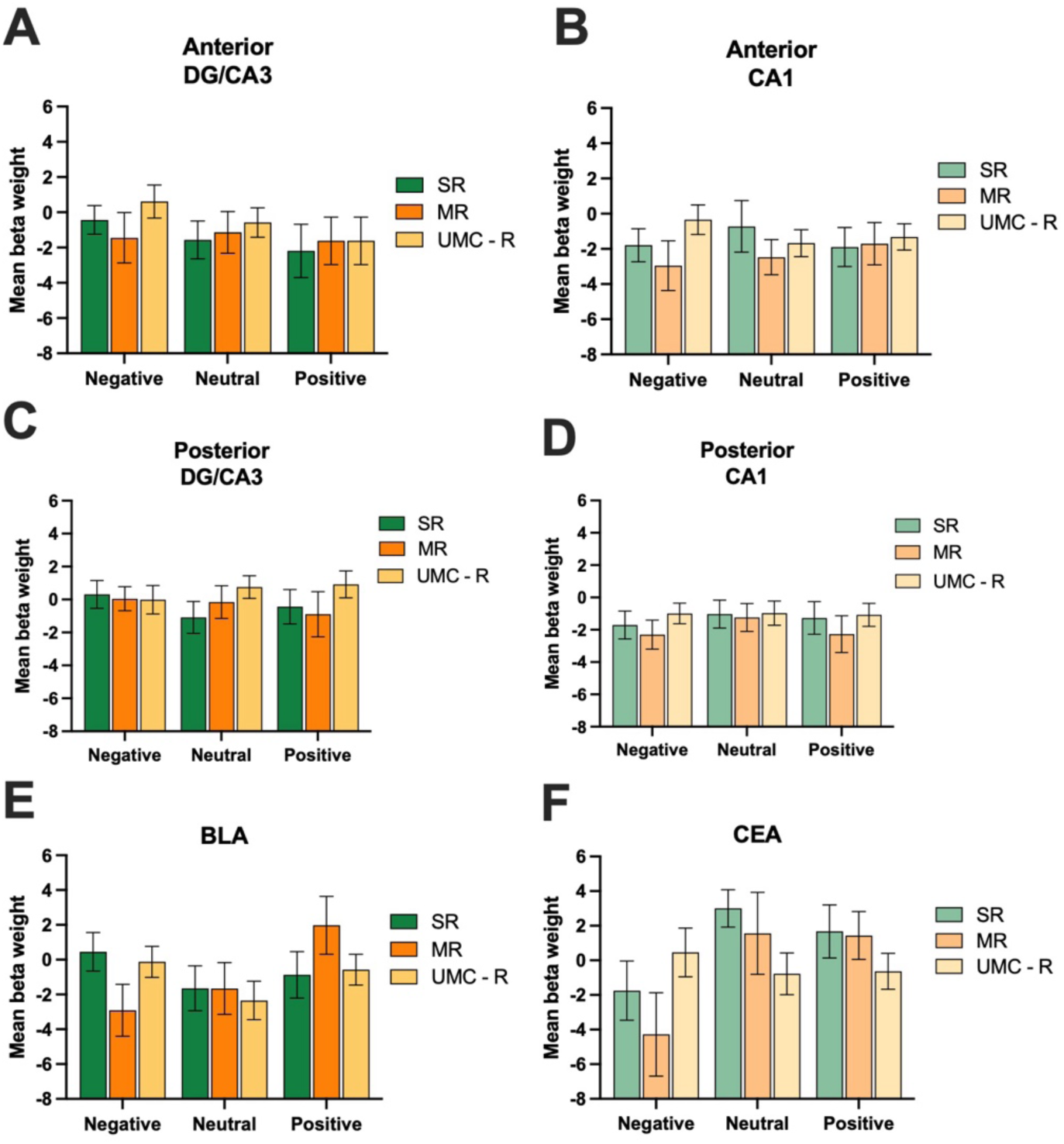
Hippocampal and amygdala activity by mechanism in R and UMC-R. A) Mean activity beta weights in the anterior DG/CA3 during lure CRs. B) Mean activity beta weights in the anterior CA1 during lure CRs. C) Mean activity beta weights in the posterior DG/CA3 during lure CRs. D) Mean activity beta weights in the posterior CA1 during lure CRs. E) Mean activity beta weights in the BLA during lure CRs. F) Mean activity beta weights in the CEA during lure CRs (*p < .05).

Next, we conducted a mixed ANOVA on lure CR activity with emotion (Negative, Neutral, Positive) and region (DG/CA3, CA1) as within-subject factors, and group (R, UMC-R) as a between-subjects factor in posterior hippocampus. There was a main effect of region [*F*(1, 146) = 14.16, *p* < .001, 17^2^ = 0.020], where there was greater DG/CA3 activity relative to CA1 (Fig. S13), but no main effects or interactions with group. When stratified by mechanism, we found a main effect of region [*F*(1, 144) = 11.82, *p* < .001, 17^2^ = 0.017], and no other main effects or interactions (Fig. S12C-D). When including hippocampal axis (anterior, posterior), we found the same main effect of region [*F*(1, 73) = 24.06, *p* < .001, 17^2^ = 0.012] in the R compared to UMC-R, as well as when stratified by mechanism [*F*(1, 72) = 18.62, *p* < .001, 17^2^ = 0.009], with no significant effects involving group or axis.

**Figure S13.**
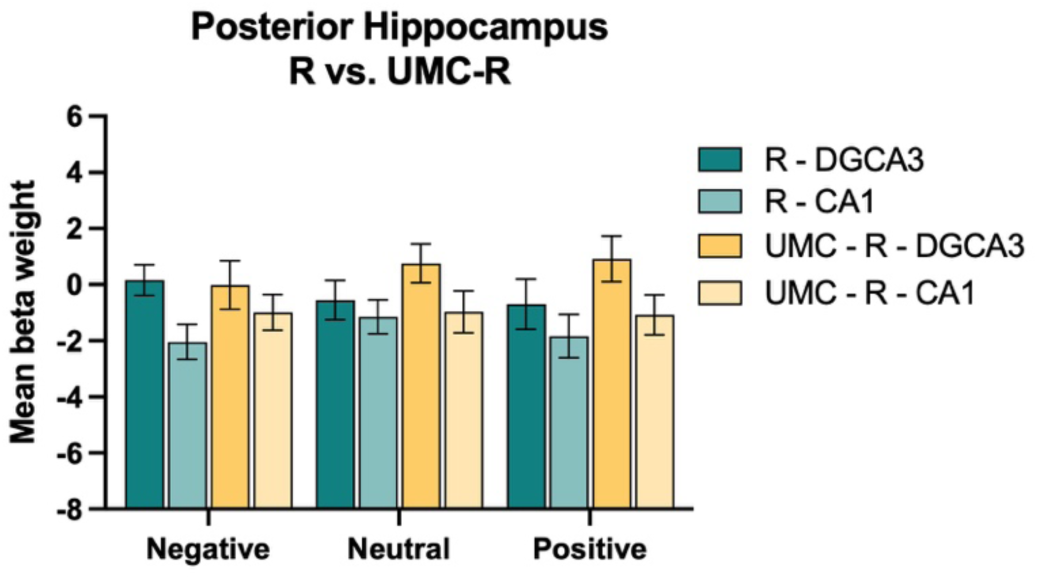
Posterior hippocampal activity in R and UMC-R. Mean activity beta weights during lure CRs (*p < .05).

Additionally, we ran the same exploratory analysis in the amygdala and stratified by mechanism. We conducted a mixed ANOVA on lure CR activity with emotion (Negative,

Neutral, Positive) and region (BLA, CEA) as within-subject factors, and group (single-action R, multi-action R, UMC-R) as a between-subjects factor. We found an interaction between emotion and region [*F*(2, 154) = 11.74, *p* < .001, 17^2^ = 0.02], where there was greater activity during neutral lure CRs in the CEA compared to the BLA but not within emotional lure CRs [*F*(1,77) = 24.60, *p* < .001, 17^2^ = 0.242] (Fig. S12E-F). There was a marginally significant interaction between emotion and group [*F*(4, 142) = 2.31, *p* = .066, 17^2^ = 0.02], where those taking multi-action antidepressants showed reduced amygdala activity during negative lure CRs and increased amygdala activity during positive lure CRs relative to both single-action and UMC-R groups [*F*(2,77) = 3.20, *p* = .046, 17^2^ = 0.077]. There was also a marginally significant three-way interaction between emotion, region, and group [*F*(4, 154) = 2.17, *p* = .075, 17^2^ = 0.006], where the above interaction between emotion and group appears to be most evident in the BLA relative to CEA [*F*(2,77) = 3.79, *p* = .027, 17^2^ = 0.090].

### Supplemental Results 8

**Figure S14.**
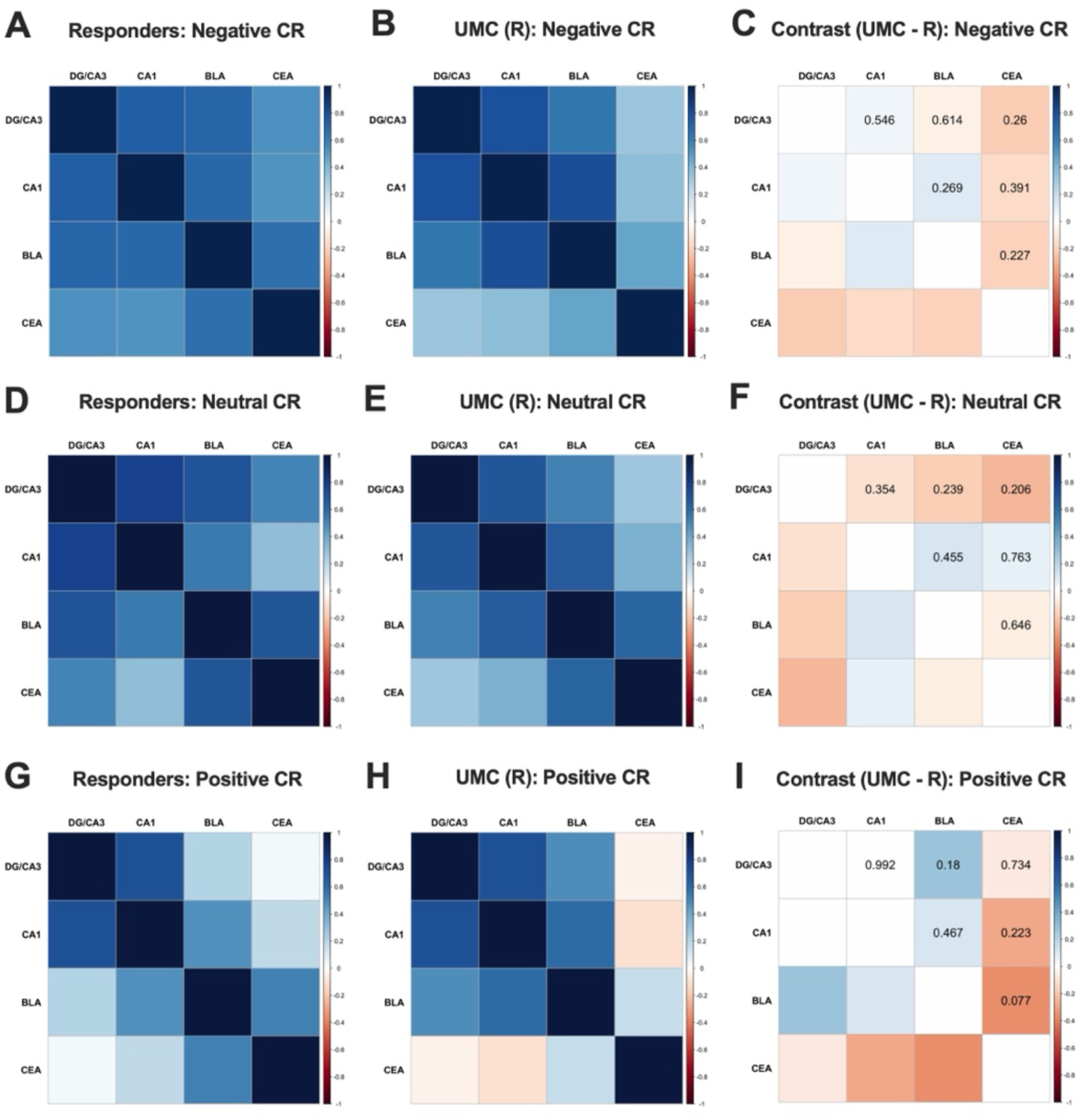
Functional coactiviation for lure CRs between R and UMC - R. A) Negative lure CRs correlation matrices for pairwise r values across ROIs in R and B) UMC-R. C) Negative lure CRs contrast correlation matrix representing the difference in pairwise r values between R and UMC-R. D) Neutral lure CRs correlation matrices for pairwise r values across ROIs in R and E) UMC-R. F) Neutral lure CRs contrast correlation matrix representing the difference in pairwise r values in R and UMC-R. G) Positive CRs correlation matrices for pairwise r values across ROIs in R and H) UMC-R. I) Positive lure CRs contrast correlation matrix representing the difference in pairwise r values in R and UMC-R. The contrast correlation matrix was normalized using Fisher’s r-to-z transformation and compared using a z-difference test (*p < .0083; Bonferroni corrected for multiple comparisons). Abbreviations: ROI, region of interest.

### Supplemental Results 9

#### No differences in hippocampal or amygdala activity between non-responders and unmedicated severity-matched controls

Next, we compared NR to UMC-NR matched on current depression severity (e.g. untreated depression). In the hippocampus, we conducted a mixed ANOVA on lure CR activity with emotion (Negative, Neutral, Positive) and region (DG/CA3, CA1) as within-subject factors, and group (NR, UMC-NR) as a between-subjects factor. There was a significant main effect of region [*F*(1, 32) = 5.86, *p* = .021, 17^2^ = 0.013], where there was greater DG/CA3 activity during lure CRs relative to CA1 (Fig. S15A), but no other significant main effects or interactions with group. This suggests that NR showed similar hippocampal activity to those who were untreated for depression (e.g. no effects of group).

**Figure S15.**
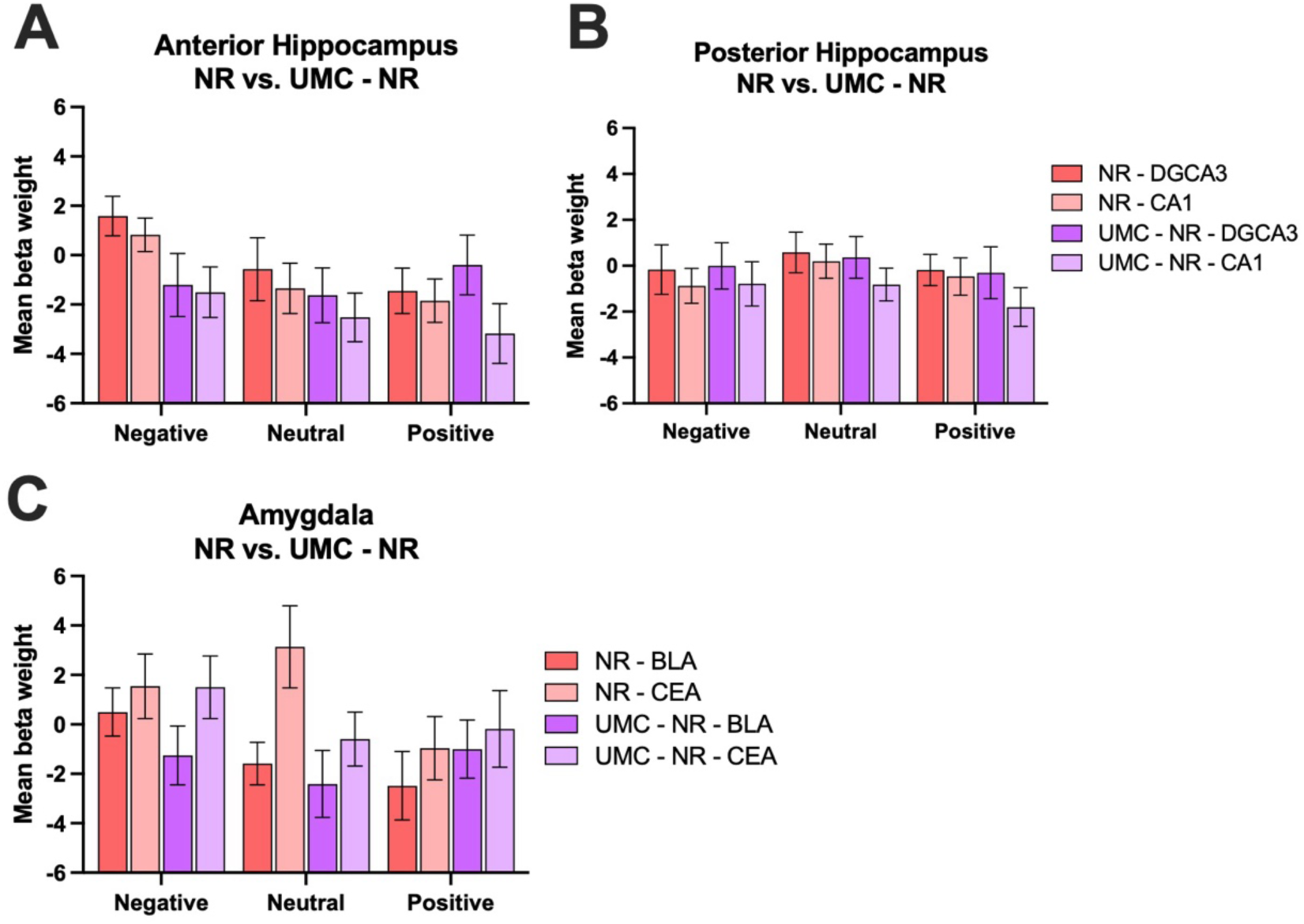
Hippocampal and amygdala activity in NR and UMC-NR. A) Anterior hippocampal mean activity beta weights during lure CRs across NR and UMC-NR groups. B) Posterior hippocampal mean activity beta weights during lure CRs across NR and UMC-NR groups. C) Amygdala mean activity beta weights during lure CRs across NR and UMC-NR groups (*p < .05).

We next conducted the same set of analyses in the posterior hippocampus. We conducted a mixed ANOVA on lure CR activity with emotion (Negative, Neutral, Positive) and region (DG/CA3, CA1) as within-subject factors, and group (NR, UMC-NR) as a between-subjects factor. We found a main effect of region [*F*(1,123) = 20.84, *p* < .001, 17^2^ = 0.030], such that there was greater activity in the posterior DG/CA3 compared to the CA1. Extending these models to include hippocampal axis (anterior, posterior), we found a main effect of region [*F*(1,32) = 7.39, *p* = .010, η^2^ = 0.012], as well as an emotion x axis interaction [*F*(2,64) = 3.82, *p* = .027, 17^2^ = 0.011]. Here, there was more activity in the posterior hippocampal regions during neutral and positive lure CRs compared to anterior hippocampal regions [*F*(1,32) = 6.73, *p* = .014, 17^2^ = 0.174] (Fig. S15B). In the amygdala, we conducted a mixed ANOVA on lure CR activity with emotion (Negative, Neutral, Positive) and region (BLA, CEA) as within-subject factors, and group (NR, UMC-NR) as a between-subjects factor. There was a significant main effect of region [*F*(1, 32) = 4.61, *p* = .039, 17^2^ = 0.017], where there was greater CEA activity during lure CRs relative to BLA (Fig. S15C), but no other significant main effects or interactions with group. This suggests that NR showed similar amygdala activity to those who were untreated for depression (e.g. no effects of group). Finally, we examined whether hippocampal-amygdala functional coactivation differed between NR and UMC-NR. There were no significant differences in hippocampal-amygdala functional coactivation across groups (Fig.S16).

**Figure S16.**
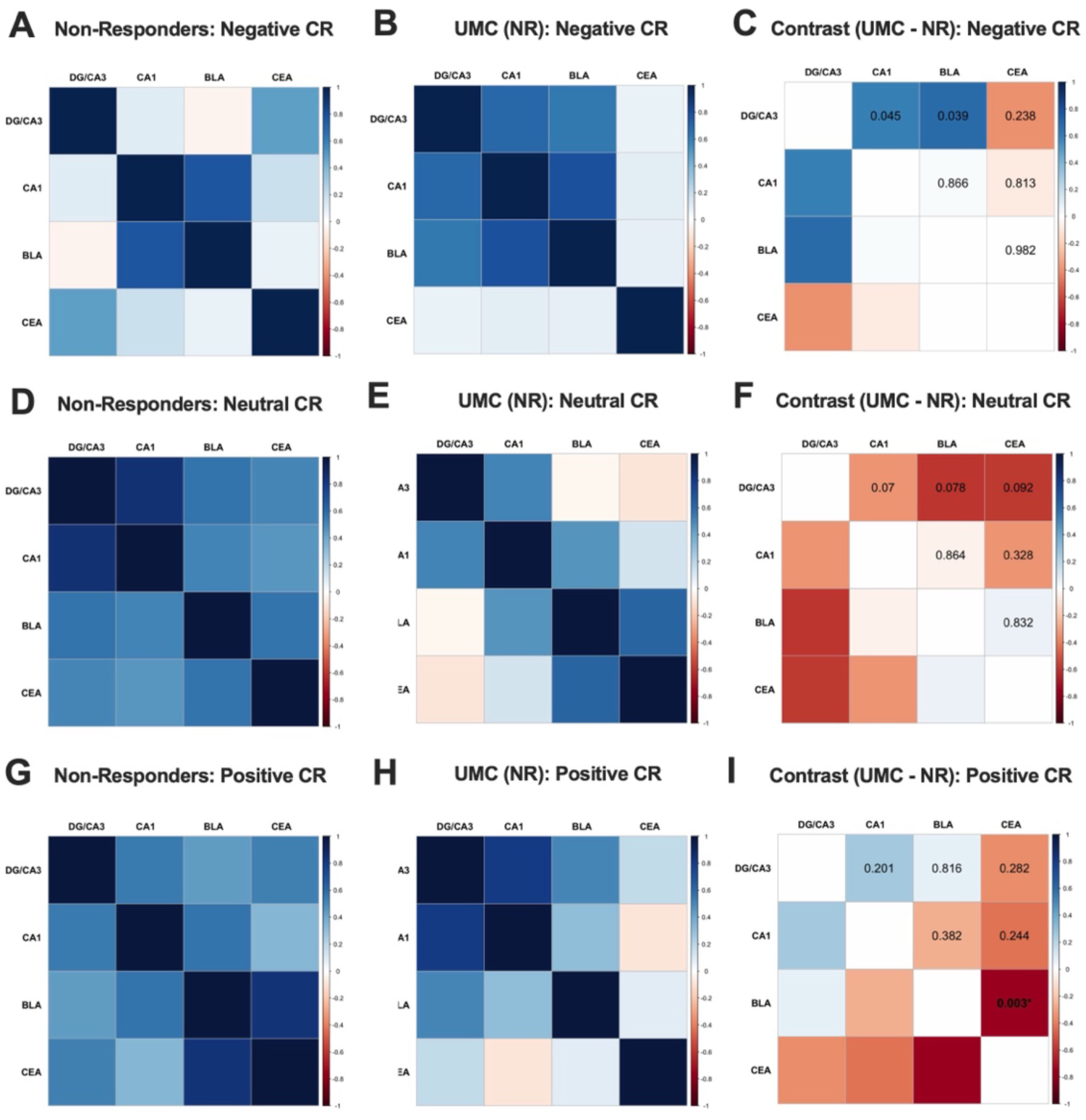
Functional connectivity for lure CRs between NR and UMC - NR. A) Negative lure CRs correlation matrices for pairwise r values across ROIs in NR and B) UMC-NR. C) Negative lure CRs contrast correlation matrix representing the difference in pairwise r values in NR and UMC-NR. D) Neutral lure CRs correlation matrices for pairwise r values across ROIs in NR and E) UMC-NR. F) Neutral lure CRs contrast correlation matrix representing the difference in pairwise r values in NR and UMC-NR. G) Positive lure CRs correlation matrices for pairwise r values across ROIs in NR and H) UMC-NR. I) Positive lure CRs contrast correlation matrix representing the difference in pairwise r values in NR and UMC-NR. The contrast correlation matrix was normalized using Fisher’s r-to-z transformation and compared using a z-difference test (*p < .0083; Bonferroni corrected for multiple comparisons). Abbreviations: ROI, region of interest.

### Supplemental Results 10

**Figure S17.**
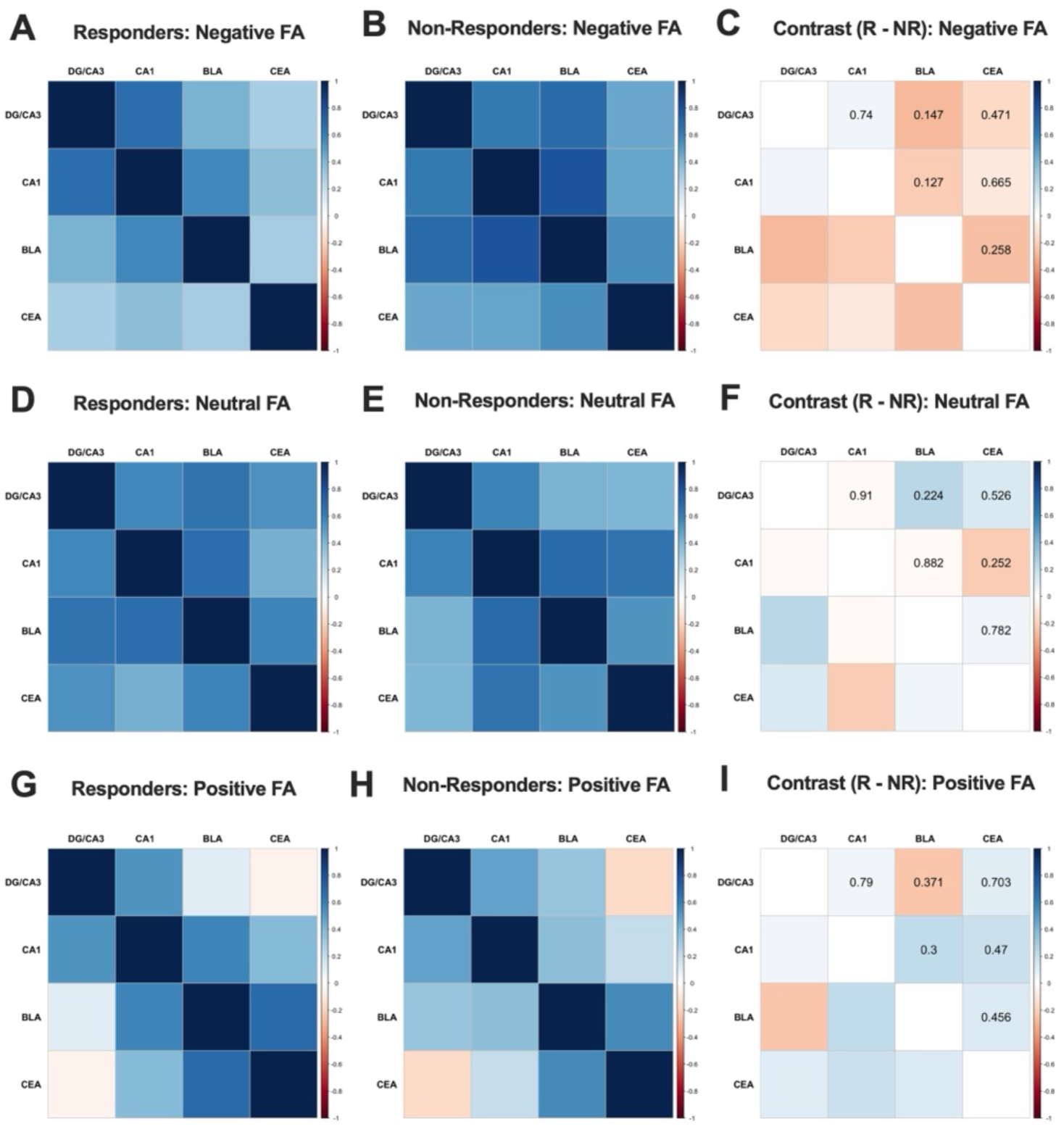
Functional connectivity for negative, neutral and positive lure FAs based on responder status. A) Negative correlation matrices for pairwise r values across ROIs in R and B) NR groups. C) Negative contrast correlation matrix representing the difference in pairwise r values in R and NR groups. D) Neutral correlation matrices for pairwise r values across ROIs in R and E) NR groups. F) Neutral contrast correlation matrix representing the difference in pairwise r values in R and NR groups. G) Positive correlation matrices for pairwise r values across ROIs in R and H) NR groups. I) Positive contrast correlation matrix representing the difference in pairwise r values in R and NR groups. The contrast correlation matrix was normalized using Fisher’s r-to-z transformation and compared using a z-difference test (*p < .0083; corrected for multiple comparisons). Abbreviations: ROI, region of interest.

**Figure S18.**
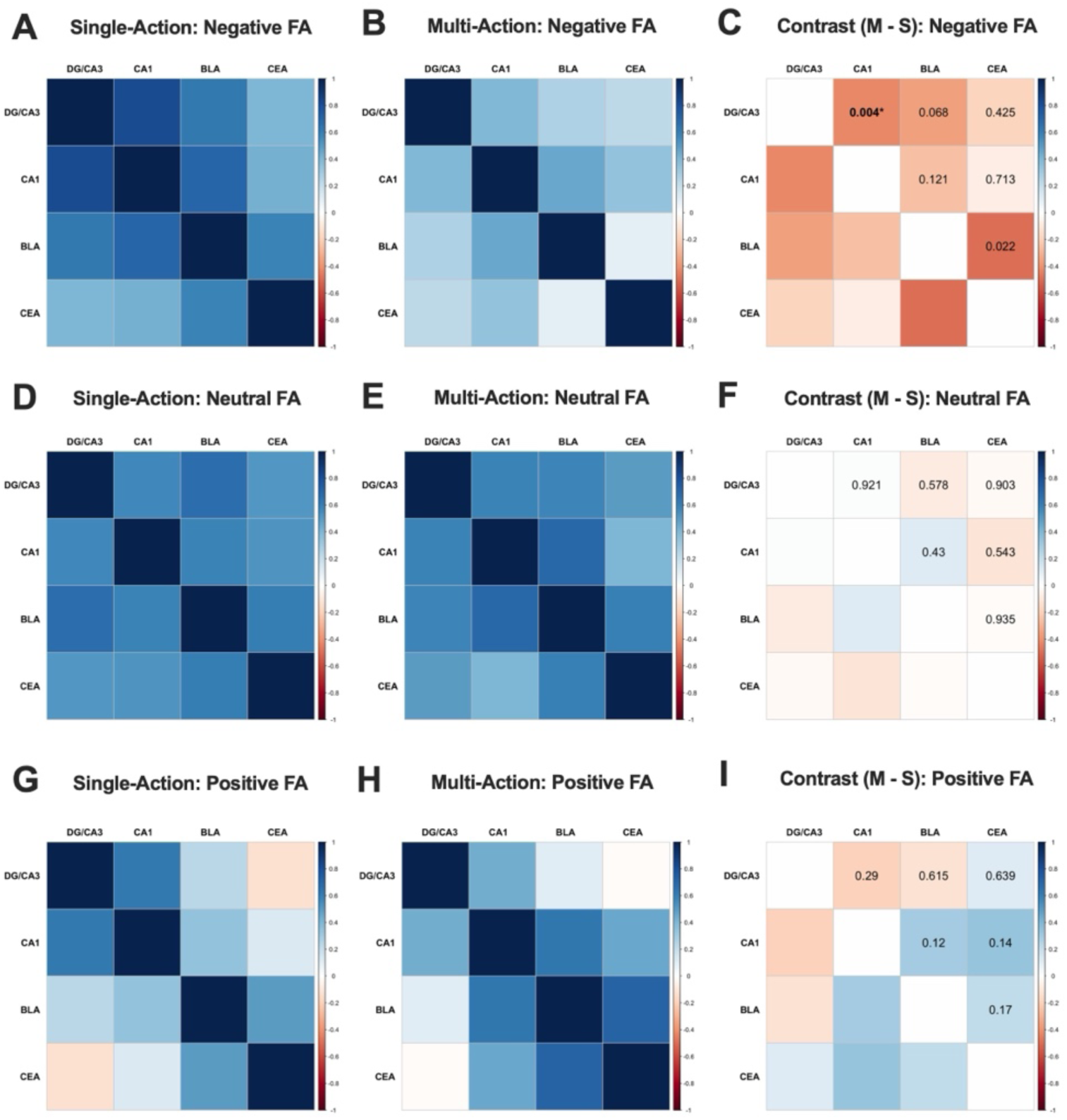
Functional connectivity for negative, neutral and positive lure FAs based on mechanism. A) Negative correlation matrices for pairwise r values across ROIs in single-action and B) multi-action groups. C) Negative contrast correlation matrix representing the difference in pairwise r values between single-action and multi-action. D) Neutral correlation matrices for pairwise r values across ROIs in single-action and E) multi-action. F) Neutral contrast correlation matrix representing the difference in pairwise r values between single-action and multi-action. G) Positive correlation matrices for pairwise r values across ROIs in single-action and H) multi-action. I) Positive contrast correlation matrix representing the difference in pairwise r values between single-action and multi-action. The contrast correlation matrix was normalized using Fisher’s r-to-z transformation and compared using a z-difference test (*p < .0083; Bonferroni corrected for multiple comparisons). Abbreviations: ROI, region of interest.

**Figure S19.**
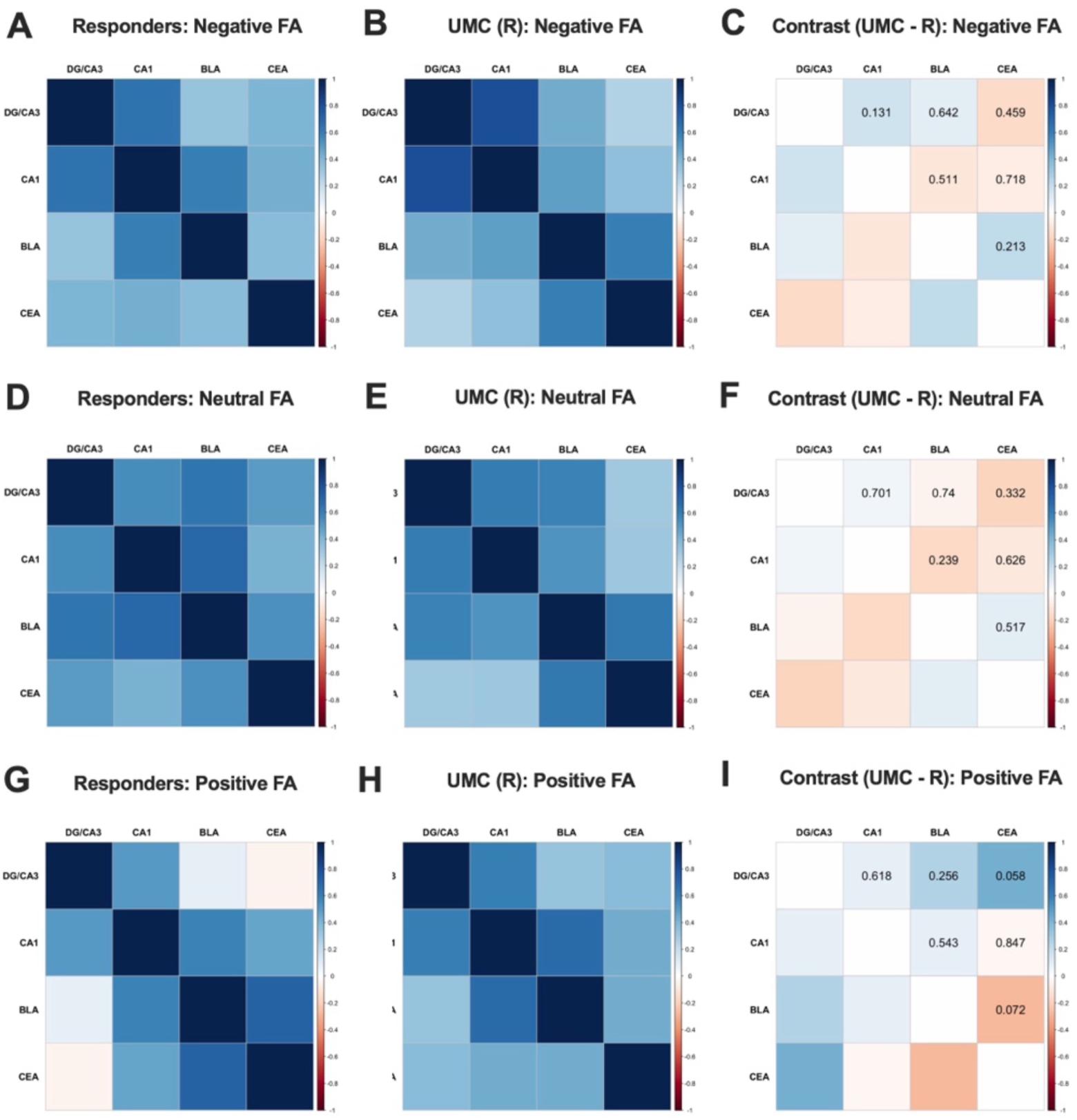
Functional connectivity for negative, neutral and positive lure FAs based in R and UMC-R. A) Negative correlation matrices for pairwise r values across ROIs in R and B) UMC-R. C) Negative contrast correlation matrix representing the difference in pairwise r values in R and UMC-R. D) Neutral correlation matrices for pairwise r^2^ values across ROIs in R and E) UMC-R. F) Neutral contrast correlation matrix representing the difference in pairwise r values in R and UMC-R. G) Positive correlation matrices for pairwise r values across ROIs in R and H) UMC-R. I) Positive contrast correlation matrix representing the difference in pairwise r values in R and UMC-R. The contrast correlation matrix was normalized using Fisher’s r-to-z transformation and compared using a z-difference test (*p < .0083; Bonferroni corrected for multiple comparisons). Abbreviations: ROI, region of interest.

**Figure S20.**
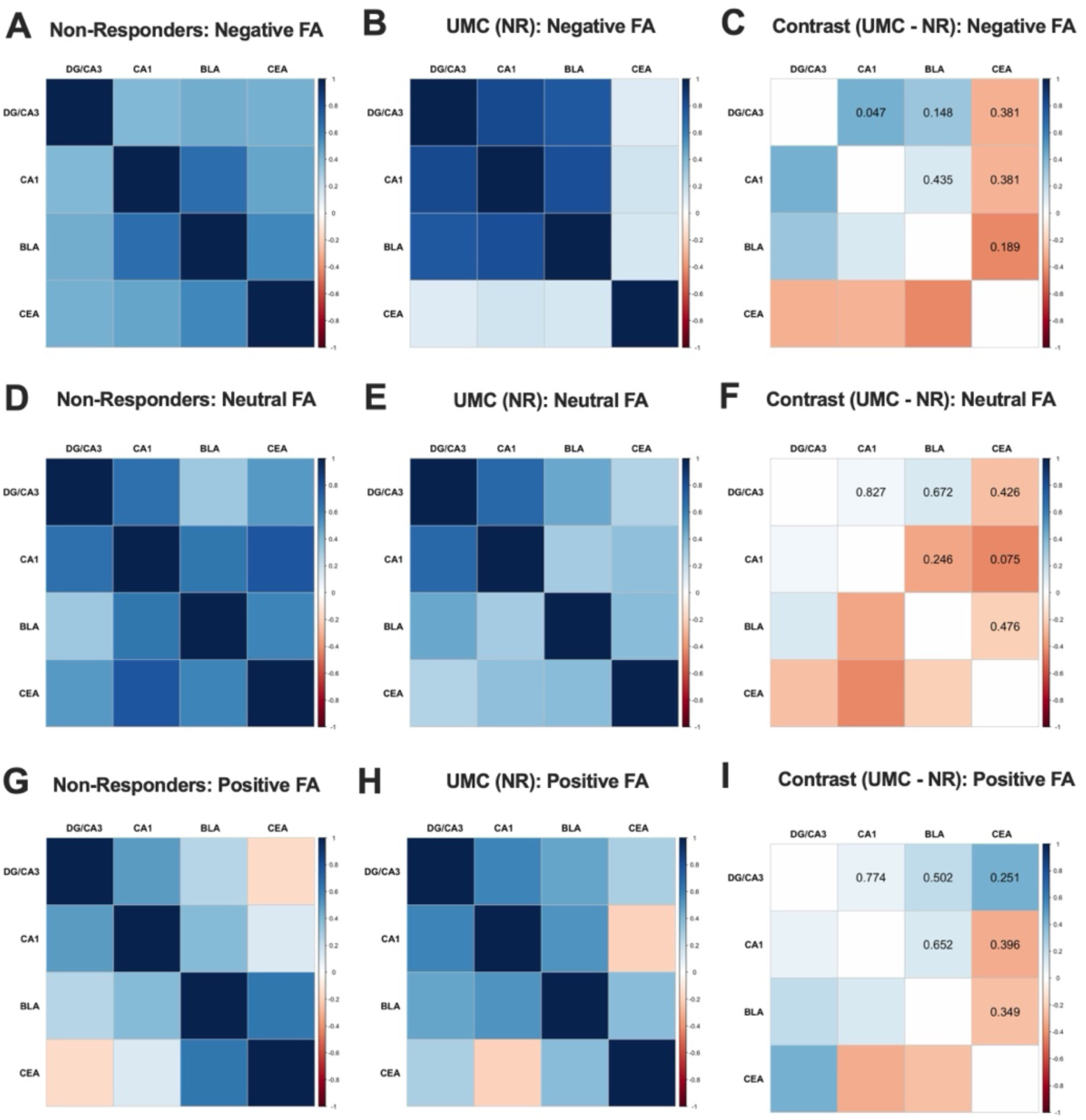
Functional connectivity for negative, neutral and positive lure FAs in NR and UMC-NR. A) Negative correlation matrices for pairwise r values across ROIs in NR and B) UMC-NR. C) Negative contrast correlation matrix representing the difference in pairwise r values in NR and UMC-NR. D) Neutral correlation matrices for pairwise r values across ROIs in NR and E) UMC-NR. F) Neutral contrast correlation matrix representing the difference in pairwise r values in NR and UMC-NR. G) Positive correlation matrices for pairwise r values across ROIs in NR and H) UMC-NR. I) Positive contrast correlation matrix representing the difference in pairwise r values in NR and UMC-NR. The contrast correlation matrix was normalized using Fisher’s r-to-z transformation and compared using a z-difference test (*p < .0083; Bonferroni corrected for multiple comparisons). Abbreviations: ROI, region of interest

